# Hox-logic of body plan innovations for social symbiosis in rove beetles

**DOI:** 10.1101/198945

**Authors:** Joseph Parker, K. Taro Eldredge, Isaiah M. Thomas, Rory Coleman, Steven R. Davis

## Abstract

How symbiotic lifestyles evolve from free-living ecologies is poorly understood. In Metazoa’s largest family, Staphylinidae (rove beetles), numerous lineages have evolved obligate behavioral symbioses with ants or termites. Widespread convergence of this lifestyle is thought to stem from a novel, chemically defended body plan that enables free-living species to infiltrate colonies and undergo extreme evolutionary specialization. Here we show how this innovative body plan evolved, via new *Hox* functions in staphylinids that remodeled the coleopteran groundplan. Using a model staphylinid, *Dalotia coriaria*, we reveal the *Hox* basis for changes in thoracic appendage morphology that shortened the beetle elytron and left the abdomen physically unprotected, selecting for an abdominal defense gland that was likely key to unlocking ant and termite societies. We present evidence that the gland evolved through a novel, combinatorial role of the abdominal Hox proteins AbdA and AbdB. These proteins function together to specify distinct gland cell types in neighboring segmental compartments, each cell type synthesizing a different class of compound—irritant, ester and solvent. Only when secreted together do these compounds constitute a bioactive secretion, providing an example of emergent chemical functionality that arises from synergy between individual gland cell types. Hox-controlled specification of glandular diversity implies a modularity in compound biosynthesis that likely catalyzed the evolvability of rove beetle chemistry, including the capacity of symbiotic taxa to produce potent compounds for host deception. This study reveals how Hox-controlled body axis modifications predispose a major animal to convergently evolve into symbionts.

---

Symbiotic relationships pervade the natural world, but their evolution from a free-living existence is largely mysterious. Explaining how a given symbiosis evolved and took on its precise form rests on explaining how the often intricate traits that mediate the relationship emerged developmentally and evolutionarily. In the Metazoa, numerous parasitic and mutualistic taxa bear specialized anatomical, physiological and behavioral adaptations for engaging in interspecies interactions. Such multifarious traits are typically idiosyncratic, lineage-specific features: the ant-tended nectary organs of lycaenid butterfly caterpillars (Hojo, Pierce, & Tsuji, 2015), the sucking disks of remoras (Britz & Johnson, 2012), or the neural differentiation of host and conspecific chatter by parasitic cowbirds (Lynch *et al*., 2017). Inferring the origins of such features can be challenging, with many appearing as novelties, or deriving from complex or extreme phenotypic modifications that cloud their evolutionary histories. Preadaptations—genetic or phenotypic attributes that evolved prior to the symbiosis itself (also termed “exaptations”: Gould & Vrba, 1982)—have proven useful for understanding the evolutionary starting material for functional traits in a variety of symbiotic relationships (Delaux et al., 2015; Henrik & Schiøtt, 2013; Parker, 2016). Preadaptive traits may form the basis for rudimentary or facultative symbioses by predisposing interactions to occur between free-living species (so called “primary preadaptations;” Parker, 2016). Preadaptations may also offer paths of least resistance to subsequent adaptation, biasing phenotypic change to certain preexisting traits as the rudimentary symbiosis evolves in intimacy (“secondary preadaptations”).

One clade that serves as a paradigm for understanding the evolution of animal symbioses are the rove beetles (Staphylinidae), currently recognized as the most species rich family in the Metazoa with >63,000 described species (Newton, 2015). Most staphylinids are free-living, predatory inhabitants of litter and soil (Grimaldi & Engel, 2005; Thayer, 2005), but numerous independent lineages have evolved to live as symbionts inside social insect colonies, in particular those of ants (myrmecophiles) and termites (termitophiles) (Kistner, 1979; 1982; Parker, 2017; Seevers, 1957; 1965). Such taxa appear to behave primarily as social parasites: burdensome colony guests, which probably impose a cost on their hosts through resource exploitation and brood predation (Akre & Rettenmeyer, 1966; Kistner, 1979; Parker, 2016; Parmentier, Bouillon, Dekoninck, & Wenseleers, 2015). The ecologies of these symbiotic species vary markedly, from opportunistic nest intruders that are attacked when detected by hosts, to socially integrated species that are accepted as apparent nestmates (Danoff-Burg, 1996; Hölldobler, 1967; 1970; Kistner, 1993; Kistner & Jacobson, 1990; Parker & Grimaldi, 2014). The repeated transition towards this form of symbiosis involves changes in anatomy and chemical ecology, as well as the evolution of novel behaviors, including complex social interactions that enable the most highly integrated species to gain acceptance inside nests.

Independent evolutionary origins of staphylinid social symbiosis are non-randomly distributed across the family, with most occurring in the Aleocharinae, the largest subfamily of ~16,000 described species (Parker, 2016; Seevers, 1957; 1965; Yamamoto, Maruyama, & Parker, 2016). The myriad instances of aleocharine myrmecophily and termitophily have been posited to stem from a preadaptive body plan. which predisposes free-living species to evolve a socially parasitic lifestyle (Parker, 2016; Seevers, 1978). Like most staphylinids, aleocharines have shortened elytra that expose an elongate, flexible abdomen (Fig 1A, B)—a morphology that permits rapid movement through soil and litter (Hammond, 1979; Hansen, 1997; Newton & Thayer, 1995). To defend the soft, otherwise vulnerable abdomen, aleocharines have evolved an exocrine “tergal” gland, which can be accurately targeted at aggressors by flexing the abdomen (Francke & Dettner, 2005; Jordan, 1913; Steidle & Dettner, 1993; Thayer, 2005). In most aleocharines, benzoquinones comprise the gland’s active volatile component (Steidle & Dettner, 1993). Benzoquinones function as effective ant deterrents (Brand, Blum, Fales, & Pasteels, 1973), enabling even free-living species to withstand worker ant aggression (Donisthorpe, 1927). The tergal gland has consequently been proposed as a primary preadaptation for social insect symbiosis, enabling aleocharines to facultatively enter nests, chemically defending themselves (Parker, 2016).

**Figure 1.**
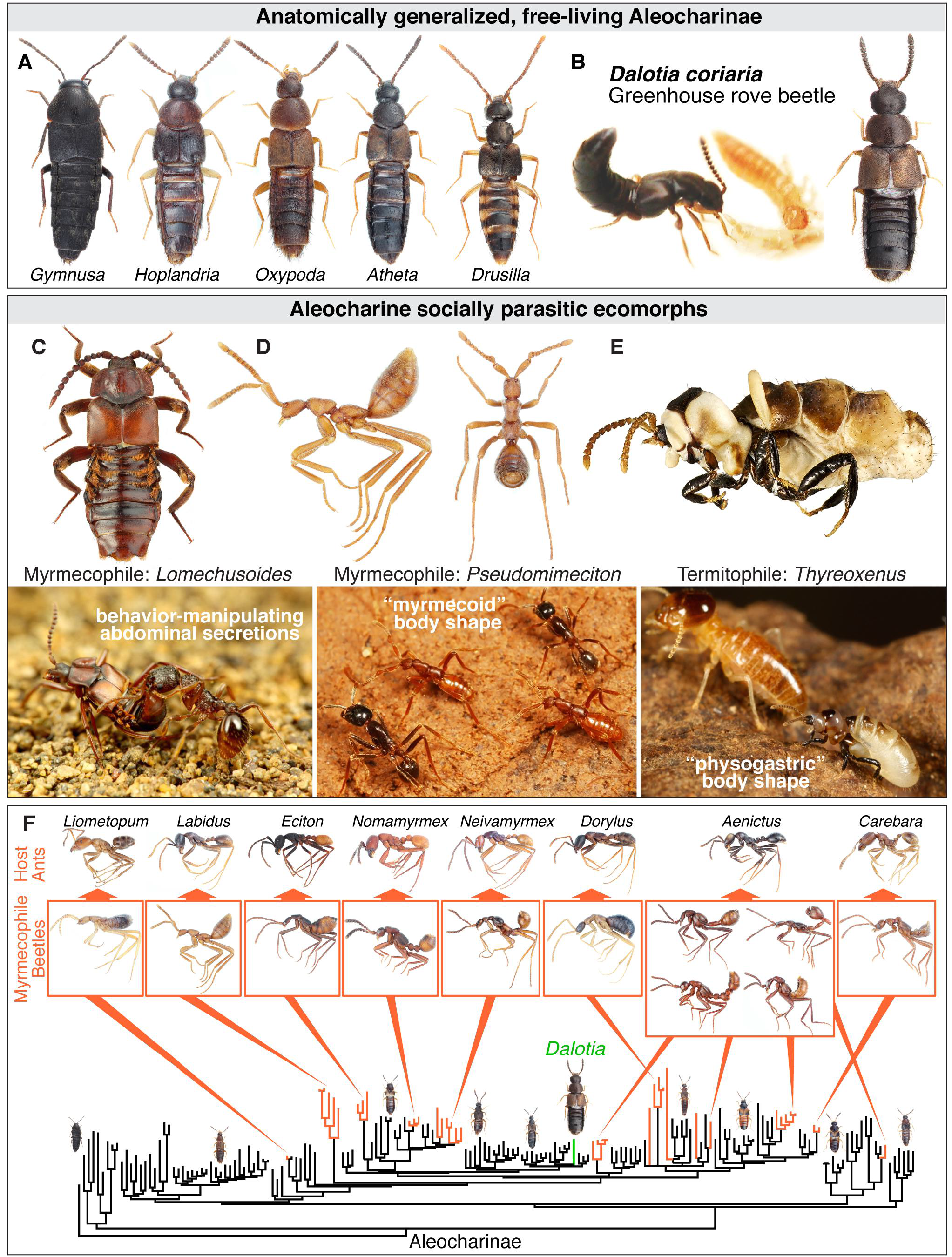
Social symbiosis in Aleocharinae. Aleocharine rove beetles are a model clade for the evolution of complex symbioses. **A**: Free living aleocharines with generalized morphology: *Gymnusa* (Gymnusini), *Hoplandria* (Hoplandriini), *Oxypoda* (Oxypodini), *Atheta* (Athetini), *Drusilla* (Lomechusini). **B**: The model aleocharine *Dalotia coriaria* Kraatz (the Greenhouse rove beetle): left, adult and larva feeding on a fly larval right, adult habitus. **C**: *Lomechusoides* myrmecophile with abdominal glands for host ant manipulation; credit: M. Smirnov (*Myrmica* ant feeding on secretion of a related *Lomechusa* beetle in Japan is shown below; Credit: T. Shimada). **D**: *Pseudomimeciton* myrmecophile with ant-like “myrmecoid” morphology (below, two living beetles walk with *Labidus* host army ants in Peru; Credit: T. Shimada). **E**: *Thyreoxenus* termitophile with termite-like “physogastric” morphology (below, *Thyreoxenus* beetle pictured with *Nasutitermes* host termite; Credit: T. Komatsu). **F**: Convergent evolution of myrmecoid syndrome in Aleocharinae: orange clades are independent origins of myrmecoid beetles, each obligately symbiotic with a different army ant genus (ants shown along the top). Representative aleocharines with generalized, ancestral morphology are shown along the bottom. The position of *Dalotia* within this convergent system is indicated, emphasizing its possession of the free-living groundplan in Aleocharinae, which has been preadaptive for repeatedly evolving social insect symbiosis. Phylogeny modified from Maruyama and Parker (Maruyama & Parker, 2017).

Tergal glands of myrmecophiles have been shown to produce novel compounds that behaviorally manipulate hosts (Akre & Hill, 1973; Hölldobler, 1970; Stoeffler, Boettinger, Tolasch, & Steidle, 2013; Stoeffler, Maier, Tolasch, & Steidle, 2007), indicating the gland can become a target for selection in species that have evolved beyond a facultative association into obligate social parasites. There has also been widespread evolution of new, specialized abdominal glands in symbiotic species that secrete unidentified chemicals that appease or otherwise influence host behavior (Fig 1C) (Parker, 2016). In many symbiotic groups, the exposed abdomen has itself become a target for selection and remodeled into shapes that mimic host ants or termites (Parker, 2016; Seevers, 1957; 1965; 1978). Abdominal shape evolution is manifested in the remarkable convergent evolution of the ant-like “myrmecoid” body form of army-ant associated aleocharines, with a narrow waist and bulbous gaster (Fig 1D) (Maruyama & Parker, 2017; Parker, 2016; Seevers, 1965). Multiple termitophile groups display a “physogastric”, termite-like body shape, where a grotesquely swollen abdomen is produced by post-imaginal growth of the fat body, with extensive intersegmental membrane between segments expanding to accommodate the balloon-like abdominal overgrowth (Kistner, 1979;: Seevers, 1957) (Fig 1E). The function of host mimicry is unclear, but it typically evolves in socially integrated species, and presumably serves to imitate tactile nestmate recognition cues (“Wasmannian mimicry”) (Kistner, 1979; Rettenmeyer, 1970).

The exposed abdomen and tergal gland of aleocharines appear to have been key to unlocking ant and termite colonies, and provide a potential system for understanding the evolution and genetic architecture of functional traits that govern animal symbioses. These features of aleocharines have conferred an adaptive flexibility to survive inside nests and undergo subsequent ecological specialization, leading to the convergent evolution of social symbiosis across the Aleocharinae phylogeny. The innovative aleocharine body plan arose via developmental modifications of the trunk and dorsal appendages of a more ancestral beetle groundplan. In insects and other arthropods, *Hox* genes confer segmental identities along the anteroposterior body axis (Lewis, 1978; Struhl, 1982). Their expression patterns and transcriptional activities underlie major anatomical differences between taxonomic groups (Hughes & Kaufman, 2002; Pearson, Lemons, & McGinnis, 2005). Here, we have determined how thoracic and abdominal *Hox* genes specify the aleocharine body plan that has been so conducive to evolving social insect symbiosis. To do this, we exploited the model organism properties of the aleocharine *Dalotia coriaria* Kraatz (=*Atheta coriaria*) (Fig 1B). *Dalotia* is a > commercially available species that is used as an agricultural biological control agent (Carney, I Diamond, Murphy, & Marshall, 2002; Echegaray & Cloyd, 2013). *Dalotia* has life history parameters that make it amenable to laboratory culture (Echegaray & Cloyd, 2013), including a: 12-20 day generation time (depending on temperature and diet), high fecundity, and sexual dimorphism that makes genetic crosses straightforward (Klimaszewski *et al*., 2007). The species is predatory, but can be cultured on artificial diets (Birken & Cloyd, 2007). Crucially, *Dalotia* is non-symbiotic, with morphology, chemistry and behavior that are generalized and I ancestral within Aleocharinae. At a phenotypic level, *Dalotia* embodies the free-living starting conditions for the convergent evolution of social insect symbiosis in this group of beetles.

## Results

### Potent gene knockdown using larval RNAi in *Dalotia*

We transformed the Green House Rove beetle, *Dalotia coriaria*, into a tractable model in which to explore gene function in staphylinids. We optimized husbandry conditions for *Dalotia* and obtained mRNA from mixed larval, pupal and adult stages. *De novo* transcriptome assembly of Illumina RNAseq reads (Grabherr *et al*., 2011) produced 46,637 isotigs (N50 = 3,091) with 22,602 isotigs ≥ 500 bp. Using this transcriptome, we synthesized dsRNAs from template cDNAs of the *Dalotia coriaria white (Dcw*) and *vermillion (Dcver)* loci, which have been shown to control eye pigmentation in beetles (Grubbs, Haas, Beeman, & Lorenzen, 2015; Lorenzen, Brown, Denell, & Beeman, 2002). We developed an RNAi protocol based on the systemic larval RNAi method used in *Tribolium* (Linz, Clark-Hachtel, Borràs-Castells, & Tomoyasu, 2014; Tomoyasu & Denell, 2004; Tomoyasu, Arakane, Kramer, & Denell, 2009; Tomoyasu, Wheeler, & Denell, 2005). Microinjection of *Dcw* or *Dcver* dsRNA at a range of concentrations into late 3^rd^ (final) instar larvae abolished eye coloration in pupae (Fig S1A–E), confirming that systemic RNAi works effectively in *Dalotia*. RNAi has an advantage over gene knockout methods in its utility for temporally controlled knockdown at a specific ontogenetic stage. We therefore used late larval RNAi to explore the functions of *Hox* genes in the formation of *Dalotia’s* imaginal structures, which grow primarily during the prepupal and pupal phases.

### Reduced growth and morphogenetic stretching underlie elytral shortening in staphylinids

BLAST searching the *Dalotia* transcriptome recovered a full complement of expressed thoracic and abdominal *Hox* genes, with no evidence of duplications (Fig S1F). We first focused on the role of the three thoracic *Hox* genes, *Sex combs reduced* (DcScr), *Antennapedia (DcAntp*) and *Ultrabithorax (DcUbx*) in sculpting one of the key morphological innovations of staphylinids—the beetle’s short elytra. Elytral shortening has been posited to be a preadaptation for social insect symbiosis in aleocharines by uncovering the abdominal segments for their subsequent chemical and anatomical elaboration (Parker, 2016; Seevers, 1978).

The beetle elytron is a modified flight wing that has become “exoskeletalized”—strengthened and rigidified via heavy chitin and pigment deposition, and by the expression of cuticular proteins (Tomoyasu, 2017). In most Coleoptera, the elytra cover approximately the entire abdomen and are similar in size or slightly smaller than the unfolded flight wings (Crowson, 1981). However, in *Dalotia*, as is typical for both aleocharines and staphylinids in general, the elytron is only 0.13× as large as the unfolded wing (Fig 2A). The wing is flight-capable and folds underneath the elytron via an efficient, origami-like pattern of folds (Saito, Yamamoto, Maruyama, & Okabe, 2014). Short elytra probably underlie Staphylinidae’s unparalleled evolutionary radiation, generating a beetle with a body plan flexible enough for undulatory locomotion through particulate substrates (Hammond, 1979; Hansen, 1997; Newton & Thayer, 1995). Developmentally, elytral size reduction in staphylinids could stem from decreased cell proliferation of the elytron relative to the wing during the prepupal growth phase. It could also arise from a non-growth related morphogenetic difference, as the cells of the wing stretch out and become larger in surface area, akin to *Drosophila* wings undergoing eversion (Roch & Akam, 2000). To determine the relative contributions of these processes to the elytronwing size discrepancy, we measured cell densities in these serially homologous structures. We used pupae rather than adults to estimate cell densities: although each cell of the adult *Dalotia* wing produces a single microtrichium, as in *Drosophila* (Fig 2C), these structures are absent from the differentiated elytron which bears only sparse setae (Fig 2B), precluding estimation of elytral cell density. In contrast, in the mature pupa (Fig 2D), cells of both wing and elytron have ceased proliferating and have secreted chitin bearing microtrichia; in both organs the cells also remain attached to the cuticle, and can be labeled with a nuclear stain to reveal cell densities (Fig 2F, G). At this stage, the elytron has attained its correct final size, but the wing has not started stretching to its full extent, which happens post-eclosion. Cell densities in the pupal wing and elytron show no significant difference (Fig 2F-H), but the elytron is nevertheless approximately half as large as the wing at this stage (Fig 2E). It follows that differences in both growth and morphogenetic stretching account for the reduced elytron size of rove beetles. A reduced relative rate of cell proliferation accounts for the initial halving of elytron size relative to the wing, seen in the pupa (Fig 2D, E). Subsequent non-proliferative, morphogenetic expansion of the wing post-eclosion exaggerates this difference still further in the adult (Fig 2A).

**Figure 2.**
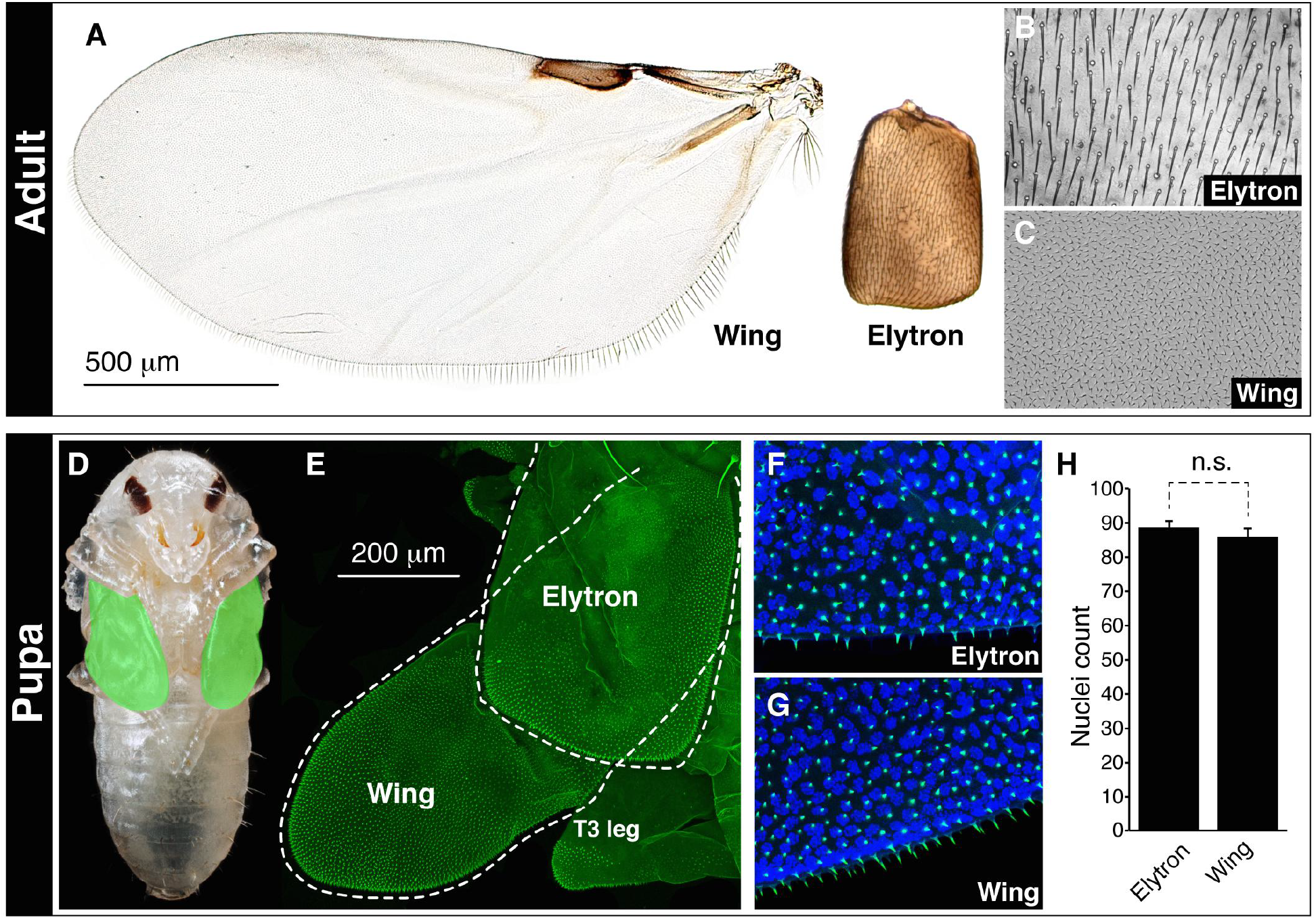
Differential growth and morphogenetic stretching underlie elytron-wing size discrepancy in rove beetles. **A:** The *Dalotia* elytron is 0.13× as large as the flight wing. **B, C**: The elytron has sparse, large setae (B), while the wing has microtrichia, each the product of a single wing cell (C). **D**: *Dalotia* pupa, ventral side, showing large flight wings (false colored in green) cloaking the body. **E**: Confocal image of cuticle autofluorescence (green) of partial *Dalotia* pupa in dorsal view, showing elytron and partially unfurled flight wing underneath, which is ~1.8× larger than the elytron at this stage. **F, G**: Confocal autofluorescence (green) and Hoechst-stained nuclei (blue) in a sector of the pupal elytron (F) and flight wing (G), showing similar cell densities in these appendages. Note that in the pupa, cells in both elytron and wing produce a microtrichium (also visible in E). **H**: Quantification of nuclear densities in elytron and wing. Error bars are S.E.M.; n.s. = not significant in a students ŕ-test.

### Hox-logic of the staphylinid short elytron

*Dalotia’s* elytron-wing size discrepancy mirrors a classical paradigm of *Hox* gene function: the transformation of the dipteran hind wing into the haltere. Here, expression of *Ultrabithorax* in the dorsal metathoracic (T3) appendage inhibits growth and morphogenetic stretching to convert the ancestral flight wing into a tiny balancing organ (Fig S2A) (Crickmore & Mann, 2006; de Navas, Garaulet, & Sanchez-Herrero, 2006; Lewis, 1978; Roch & Akam, 2000; Weatherbee, Halder, Kim, Hudson, & Carroll, 1998). Elytral size reduction in staphylinids might arise from similar Hox-dependent modulation of organ size, but with the size decrease occurring in the mesothoracic (T2) appendage and hence under the influence of a more anterior *Hox* gene. Counter to this idea, however, studies in *Drosophila* and *Tribolium* have shown that the dorsal T2 appendage—the wing or elytron, respectively—arises independently of *Hox* input, since loss of any of the three thoracic *Hox* genes does not impact its development (Struhl, 1982; Tomoyasu et al., 2005). We tested whether the staphylinid elytron also represents the default, Hox-free appendage state. Knockdown of *DcUbx* by microinjecting dsRNA into late third instar larvae induced a classical bithorax mutant phenotype in the resulting pupa (Fig 3A, B) (Lewis, 1978). The T3 appendage (the wing) converted to the T2 identity (elytron) (Fig 3A, B), and the scutellum, a T2 structure, was also duplicated in T3 (Fig 3A, B). This result confirms that *DcUbx’s* role is confined to T3, where it blocks elytron exoskeletalization to specify an enlarged, membranous flight wing, as in *Tribolium* (Tomoyasu et al., 2005). Crucially, loss of the two anterior thoracic *Hox* genes, *DcScr* and *DcAntp*, also caused negligible effects on *Dalotia’s* elytra: knockdown of *DcScr* caused the appearance of elytron-like outgrowths from the pronotum (T1), consistent with *Scr’s* role in repressing dorsal appendage development in the insect prothorax (Carroll, Weatherbee, & Langeland, 1995; Tomoyasu et al., 2005), but in T2, elytron size and morphology were not detectably affected (Fig 3C, D). Similarly, DcAntp-RNAi produced a malformed leg phenotype, without any discernable effect on the elytron (Fig 3C, E). The rove beetle elytron thus appears to represent the developmental “ground state”—the default morphology of the dorsal thoracic appendage, which develops without influence of *Hox* genes.

**Figure 3.**
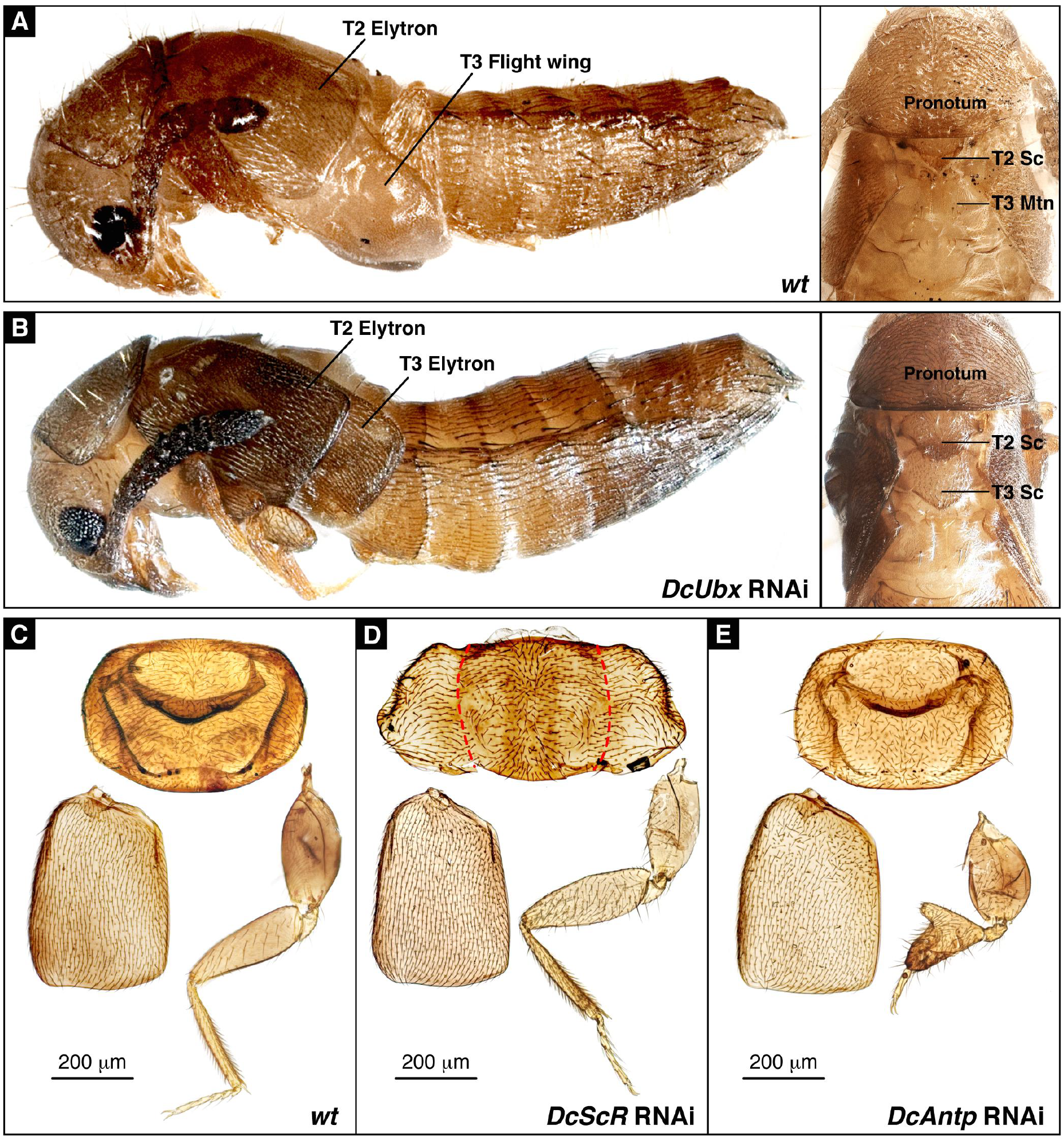
The elytron develops as a *Hox-free* appendage in rove beetles. **A**: Wild type pupa showing T2 elytron and T3 flight wing; inset shows dorsal scutellum (Sc) on segment T2 and metanotum (Mtn) on T3. **B**: *DcUbx* RNAi pupa showing the elytron and scutellum are now duplicated in T3. **C**: Wild type adult pronotum, elytron and T1 leg. **D**: *DcScr* RNAi causes elytron-like outgrowths from the edges of the pronoum, but leaves the elytron and T1 legs largely unaffected. **E**: *DcAntp* RNAi causes a malformed leg phenotype but leaves the pronotum and elytron unaffected (note that two different DsRNAs targeting different regions of the *DcAntp* transcript independently reproduced this same phenotype).

### Evolutionary novel function of rove beetle *Ubx* in promoting wing growth

The mechanism of organ size reduction in the rove beetle elytron differs to that operating in the fly haltere, where ground state size is modified by the growth-repressive function of *Ubx* (Crickmore & Mann, 2008). Rather, the staphylinid elytron shares the Hox-free ground state property with the dorsal T2 appendage of most other holometabolan insects (Struhl, 1982; Tomoyasu, 2017; Tomoyasu et al., 2005), but the ground state has itself been intrinsically reprogrammed during evolution so that the elytron attains only a small size. This staphylinid-specific mechanism has consequences for the function of *Ubx* in rove beetles. For the animal to develop enlarged hind wings capable of flight, *Ubx* must act in T3 to override the size reduction inherent to the ground state (Fig 3A, B). Staphylinid *Ubx* has thus evolved to function effectively as a growth-promoting transcription factor—a novel role within the Coleoptera, and one that is opposite to its growth-repressive activity in the dipteran haltere (Crickmore & Mann, 2006; de Navas et al., 2006; Weatherbee et al., 1998). How does DcUbx produce a different developmental output to its *Drosophila* ortholog? Radical changes in *Hox* function during evolution have arisen by both cis-regulatory changes in enhancers of Hox target genes (e.g. Jeong, Rokas, & Carroll, 2006), and also via evolution of the Hox protein sequences themselves, via acquisition of novel transcriptional activation or repression domains (Galant & Carroll, 2002; Ronshaugen, McGinnis, & McGinnis, 2002; Tour, Hittinger, & McGinnis, 2005). The opposite effects of Ubx in *Dalotia* and *Drosophila* could depend on the species-specific genomic contexts in which the proteins ordinarily function. Conversely, it could stem from divergent Ubx protein sequences, which share only 52% amino acid identity between the two species, conferring potentially opposite activities on transcription when bound to common loci.

To distinguish between these alternatives, we cloned *DcUbx* for GAL4/UAS-mediated misexpression in *Drosophila*, and compared the effects to expressing *Drosophila Ubx (DmUbx*). We used *nubbin-GAL4* to drive transgenes in the blade territory of the wing imaginal disc, temporally restricting GAL4 activity specifically to the third larval instar by using temperature sensitive GAL80 and shifting to the permissive temperature (McGuire, Le, Osborn, Matsumoto, & Davis, 2003). As shown previously (Pavlopoulos & Akam, 2011), expression of *DmUbx* using this method strongly represses growth and causes the blade to transform into a tiny haltere-like structure (Fig S2B, C). Strikingly, *DcUbx* produced a near-identical phenotype in the *Drosophila* wing (Fig S2B, D), indicating that *DmUbx* and *DcUbx* possess the same growth repressive activity when mis-expressed in flies. As an additional test we looked at the impact of the two Ubx proteins on morphogen production. In the haltere, DmUbx has been shown to repress growth in part by impeding production of the Decapentaplegic (Dpp) morphogen (Crickmore & Mann, 2006; de Navas et al., 2006). Expressing DmUbx in the Dpp expression domain of the wing reduced the output and range of Dpp, as revealed by phosphorylated MAD (pMAD), and decreased the size of the entire blade (Fig S2E, F). Using this assay, *DcUbx* again behaved similarly (Fig S2E, G). This equivalence of *DmUbx* and *DcUbx* activity in *Drosophila* implies that their opposing phenotypic effects in their native contexts are unlikely to arise from differences in protein sequence. Instead, divergent regulatory evolution downstream of Ubx, in loci tasked with overriding the ground state, accounts for why DmUbx produces a diminutive haltere from an enlarged flight wing, while DcUbx produces an enlarged flight wing from a diminutive elytron.

### Ontogeny and chemistry of the tergal gland: an evolutionary key innovation

By exposing the abdomen, the evolutionary development of short elytra imposed an “Achilles’ heel” on staphylinids, with various subfamilies countering the lack of physical protection by evolving abdominal defense glands (Araujo, 1978; Dettner, 1993; Parker, 2017; Thayer, 2005). In Aleocharinae, evolution of a tergal gland on the dorsal abdomen appears to have had major consequences for the subfamily’s evolutionary success. The vast majority of the >16,000 described species, including *Dalotia*, comprise one clade, the so-called “higher Aleocharinae”, which is defined by the presence of the tergal gland and occupies a spectrum of ecological niches that is unprecedented within the Coleoptera (Thayer, 2005; Yamamoto et al., 2016). In contrast, four earlier-diverging tribes that form the remainder of the subfamily lack the gland, and number only ~120 species (Ashe, 2005; Steidle & Dettner, 1993; Yamamoto et al., 2016). The glandless outgroup subfamilies Tachyporinae, Trichophyinae, Phloeocharinae and Habrocerinae are similarly species-poor relative to the higher Aleocharinae. Evolution of the tergal gland appears to represent a defensive “key innovation”, which helped catalyze the higher Aleocharinae’s radiation in terrestrial ecosystems.

The developmental changes in the higher aleocharine stem lineage that underlie tergal gland evolution are unknown. Likewise, the gland’s genetic capacity to synthesize defensive compounds, which can vary in a species-specific fashion to adapt the beetles to different habitats, is a mystery. In adult *Dalotia*, as in most other Aleocharinae, the tergal gland appears as a large, sack-like reservoir of transparent, chitinous intersegmental membrane extending from the anterior edge of abdominal tergite 7 (A7) (Fig 4A). The margin of tergite 7 projects outward, forming a sclerotized cuticular shelf (Fig 4A), which regulates the release of the secretion from the reservoir. When filled, the defensive secretion has a yellow color (Fig 4A), and by twisting the abdomen over the body, the gland contents can be expelled at or smeared on aggressors (Araujo, 1978; Brand et al., 1973; Parker, 2016) (Supplemental Video S1). We used gas chromatography-mass spectrometry (GC-MS) to profile *Dalotia’s* tergal gland chemistry, collecting the secretion via three different methods: dabbing filter paper onto the gland opening (Fig 4B, Fig S3A), forcing the beetle to expel the gland contents directly into hexane by briefly submersing it (Fig S3B), and using dynamic headspace collection of volatiles (Fig S3C). All three methods detected largely similar proportions of the same compounds: three benzoquinones, responsible for the secretion’s yellow color (1,4-benzoquinone, 2-methyl-1,4-benzoquinone and 2-methoxy-3-methyl-1,4-benzoquinone), two esters (ethyl decanoate and isopropyl decanoate, the latter not detected in the headspace) and large amounts of an alkane, n-undecane (Fig 4B; Fig SS3). The n-undecane functions as the hydrocarbon solvent for the benzoquinones (Steidle & Dettner, 1993), and the esters are probably surfactants, or wetting agents, facilitating the spreading of the secretion across biological tissues (Dettner, 1984).

**Figure 4.**
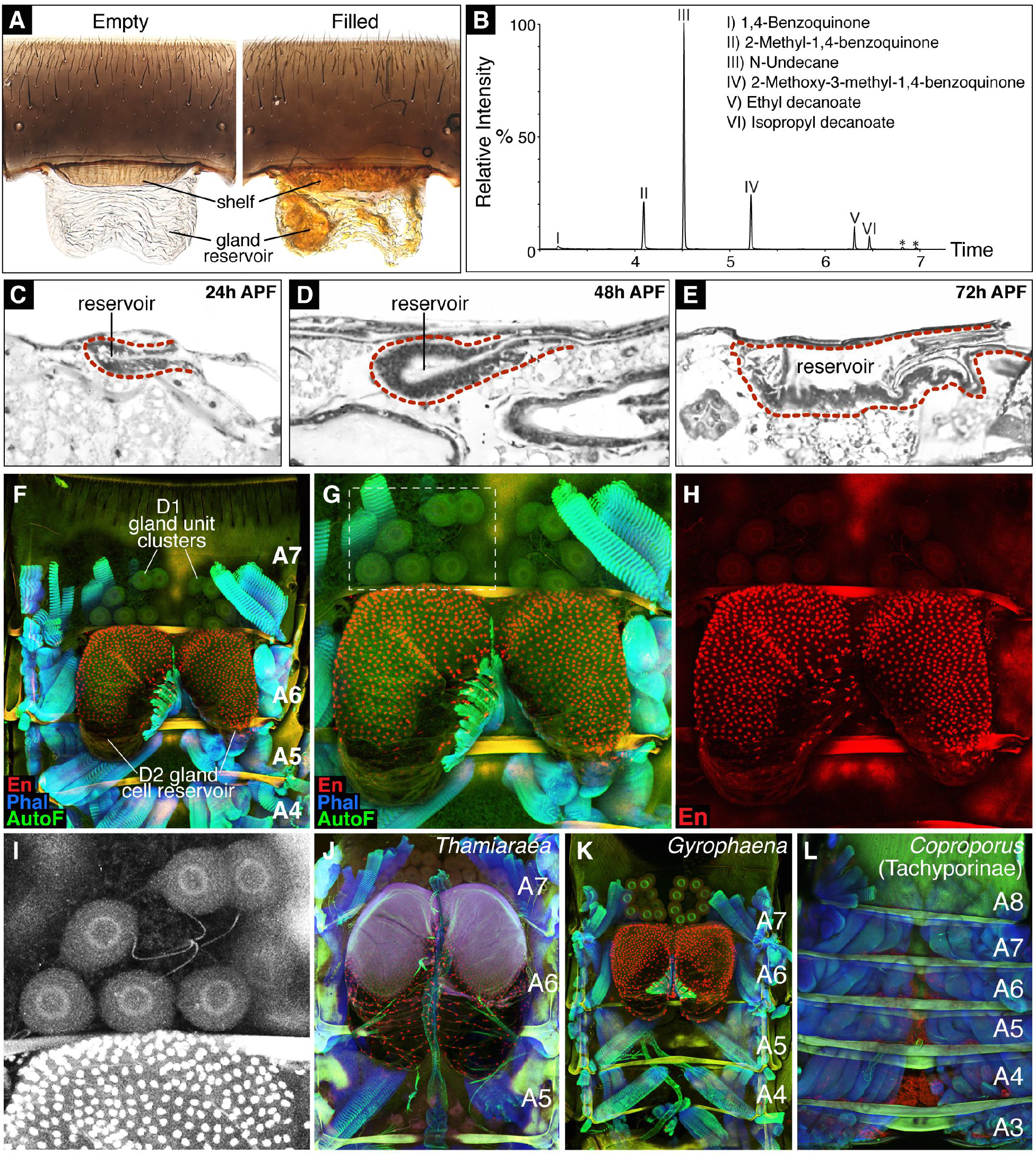
Ontogeny and defensive chemistry of the *Dalotia* tergal gland. **A**: Removal of abdominal segment A7 reveals *Dalotia’s* tergal gland reservoir: a thin bilobed sack of intersegmental membrane that contains yellow benzoquinones when filled. The anterior margin of tergite 7 is fashioned into a shelf that seals off the gland opening. **B**: GC-MS of the *Dalotia* tergal gland, with compounds corresponding to peaks listed. Asterisks denote contaminant compounds from tissue paper dabbed on the tergal gland opening; these are absent when other methods of chemical extraction are used (See Supplemental Fig S3). **C–E**: Sagital histological sections of *Dalotia* pupae from 24-96 hours after puparium formation, showing the timecourse of gland development from a small invagination of cells budding from the dorsal epidermis (C), to a large folded epithelial sack (E). **F**: Confocal image of the dorsal internal adult abdomen of *Dalotia*, labeled for En protein (red), Phalloidin-stained muscle (blue) and autofluorescence (AutoF; green). The tergal gland is prominent and sits directly underneath segments A6 and A7. The gland consists of a large, bilobed reservoir comprised of En-positive D2 secretory cells, and two clusters of large, En-negative D1 gland units. **G, H**: Enlarged region of F showing tergal gland organization; boxed region is shown further enlarged in I. En protein staining (red) shows that the D2 reservoir cells express En, and hence are of P-compartment origin from segment A6; The D1 gland unit cells do not express En, and are likely derived from the A-compartment of segment A7. I: The D1 gland units are a classical gland type consisting of a large bulb attached to a duct, which feeds into the reservoir (Araujo, 1978). **J-L**: Confocal image of the dorsal internal adult abdomen of three other staphylinids, labeled for En protein (red), Phalloidin-stained muscle (blue) and autofluorescence (green). The two aleocharines *Thamiaraea* (J) and *Gyrophaena* (K) possess a tergal gland; in contrast, *Coproporus*, of the outgroup subfamily Tachyporinae, is glandless (L).

*Dalotia’s* benzoquinone/undecane-based secretion matches the quinone/hydrocarbon chemistry of most other free-living aleocharines studied thus far (Steidle & Dettner, 1993). This general composition likely approximates the primitive tergal gland chemistry within the subfamily—one that has been augmented or substituted with novel compounds in at least some social insect symbiont taxa (Stoeffler et al., 2007; 2013; Stoeffler, Tolasch, & Steidle, 2011). We explored how the gland forms developmentally. Unlike the wing and elytron, which develop primarily during the prepupal phase and appear fully grown in the pupa, we found the tergal gland reservoir arises later, as an invaginating pocket of dorsal ectoderm during pupation (Fig 4C–E). Using histological sections, we observed the nascent reservoir as a small cluster of cells budding from the epidermis at twenty-fours hours after pupal formation (Fig 4C). The reservoir cells subsequently proliferate over the next 48 hours, producing an extensively folded columnar epithelial sack inside the body cavity (Fig 4D, E) (Araujo & Pasteels, 1985). Tergal gland reservoir development is akin to the imaginal discs of *Drosophila*, which invaginate from the ectoderm and form an internal sack of epithelial cells that proliferate inside the body cavity (Held, 2005). Unlike the imaginal discs, however, the gland reservoir never undergoes morphogenetic eversion, remaining invaginated within the abdomen.

### *Hox* logic of tergal gland development and chemical biosynthesis

Dissecting open the dorsal adult abdomen exposes the organization and anatomy of the mature gland, revealing a composite structure (Fig 4F, G). The gland reservoir is a bilobed sack that sits directly underneath A6, and associated with it are two symmetrical clusters of 6–8 large, classical gland units (Fig 4F, G). Each unit is comprised of a globular bulb cell with a large, internal extracellular space, and a duct cell (Fig 4I). These gland units are situated directly posterior to the reservoir in the anterior portion of A7 (Fig 4F, G, I). The cells comprising the reservoir are of a ductless secretory type that has been termed “D2”, while the large, classical gland units have been termed “D1” (Araujo, 1978; Araujo & Pasteels, 1985; Steidle & Dettner, 1993)—a terminology we follow here. The ducts of the D1 clusters feed into the D2 reservoir (Araujo, 1978; Steidle & Dettner, 1993), and based on their enlarged nuclei (Fig S5D, E), the large D1 bulb cells appear to be polyploid. How did this evolutionary novelty form at this specific abdominal position? Developmental patterning of the adult insect abdomen has been studied in *Drosophila*, where the intersegmental membrane between tergites derives from cells comprising the posterior (P) compartment (Kopp & Duncan, 2002; Struhl, Barbash, & Lawrence, 1997). We speculated that the D2 gland cells that comprise the reservoir, and which are continuous with the A6-A7 intersegmental membrane, might be of P compartment origin. Staining the adult gland for Engrailed (En) protein, which specifies P compartment identity (Morata & Lawrence, 1975), reveals that all D2 gland cells label strongly for En (Fig 4F–H). The reservoir thus originates from the P compartment of segment A6. In contrast, the D1 gland units sitting behind the reservoir do not label for En (Fig 4H), indicating they likely derive from the anterior (A) compartment cell population of segment A7. We explored a variety of other higher aleocharine species, and found all of them to have a glandular organization similar to that of *Dalotia* (Fig 4J, K, Fig S4A–C). Each species possesses a gland reservoir under segment A6 that is derived from Engrailed-positive posterior compartment cells, together with a varying number of D1 gland units situated directly behind, in A7 (Fig 4J, K, Fig S4A–C). In contrast, *Coproporus* from the closely allied subfamily, Tachyporinae, exhibited no such structure (Fig 4L), and the gland is also known to be absent from the early-diverging aleocharine tribes Gymnusini, Mesoporini and Trichopseniini (Ashe, 2005). The gland is thus a morphological and chemical novelty of higher Aleocharinae, which develops in a specific segmental position and with a stereotyped compartmental organization.

We explored the potential roles in gland evolution of the two *Hox* genes that function in the insect abdomen, *Abdominal A (AbdA*) and *Abdominal B (AbdB*). Using whole-mount cuticle preps of the *Dalotia* adult abdomen, we determined the regions of segmental specification that are under the respective control of the AbdA and AbdB Hox proteins (Fig S5A–C). RNAi-knockdown of *AbdA* caused a segmental transformation phenotype that extended from segment A3 all the way to the posterior margin of A7, which takes on a jagged form, similar to that of A8 (Fig S5A, B). Hence, AbdA controls segmental identities at least as far posteriorly as A6 and A7, the gland-bearing segments. Notably, in *AbdA* knockdown animals, the shelf at the anterior margin of A7, which marks the tergal gland opening in wild type animals (Fig S5A), was missing (Fig S5B), and we could also see no clear evidence of a membranous gland reservoir, implying a potential role for AbdA in tergal gland specification. In AbdB knockdown animals, all abdominal segments, including A7 and A8, assume an identity similar to A3–A6 (Fig S5A, C), and again, no cuticular shelf on A7 nor membranous reservoir on A7 could be observed. Hence, the morphology of A6 and A7, the gland bearing segments, are specified by a combination of both AbdA and AbdB acting together. Investigating the internal abdomen, we found that *AbdA* RNAi completely abolished tergal gland formation, with both the D2 reservoir and D1 gland cells failing to develop (Fig 5A, B). Nuclei of both the D1 and D2 gland cells stain positively for AbdA using the antibody FP6.87, indicating that AbdA protein is active in both cell types (Fig S5D, E; note that the epitope for FP6.87 antibody is shared with Ubx, but based on *Drosophila*, expression of this protein is only in more anterior abdominal segments (Kopp & Duncan, 2002)). Similarly, *AbdB* RNAi caused total loss of tergal gland structures (Fig 5C). We conclude that *AbdA* and *AbdB* function combinatorially and non-redundantly to specify the tergal gland. Loss of either Hox protein prevents gland formation (Fig 5A–C), effectively phenocopying the glandless condition of higher aleocharine sister lineages (Fig 4L). We therefore propose that during evolution of the higher aleocharine stem lineage, AbdA and AbdB acquired novel functions, which enabled them to act together in a region of overlapping expression that spans the interface between segments A6 and A7. In these neighboring segments, the two Hox proteins synergistically to specify the tergal gland, but they do so via distinct outputs in abutting segmental compartments (Fig 5H). In the P compartment of A6, AbdA and AbdB produce the epidermal invagination of D2 secretory cells that comprise the tergal gland reservoir. In contrast, in the A compartment of segment A7, AbdA and AbdB specify the D1 gland cells (Fig 5H).

**Figure 5.**
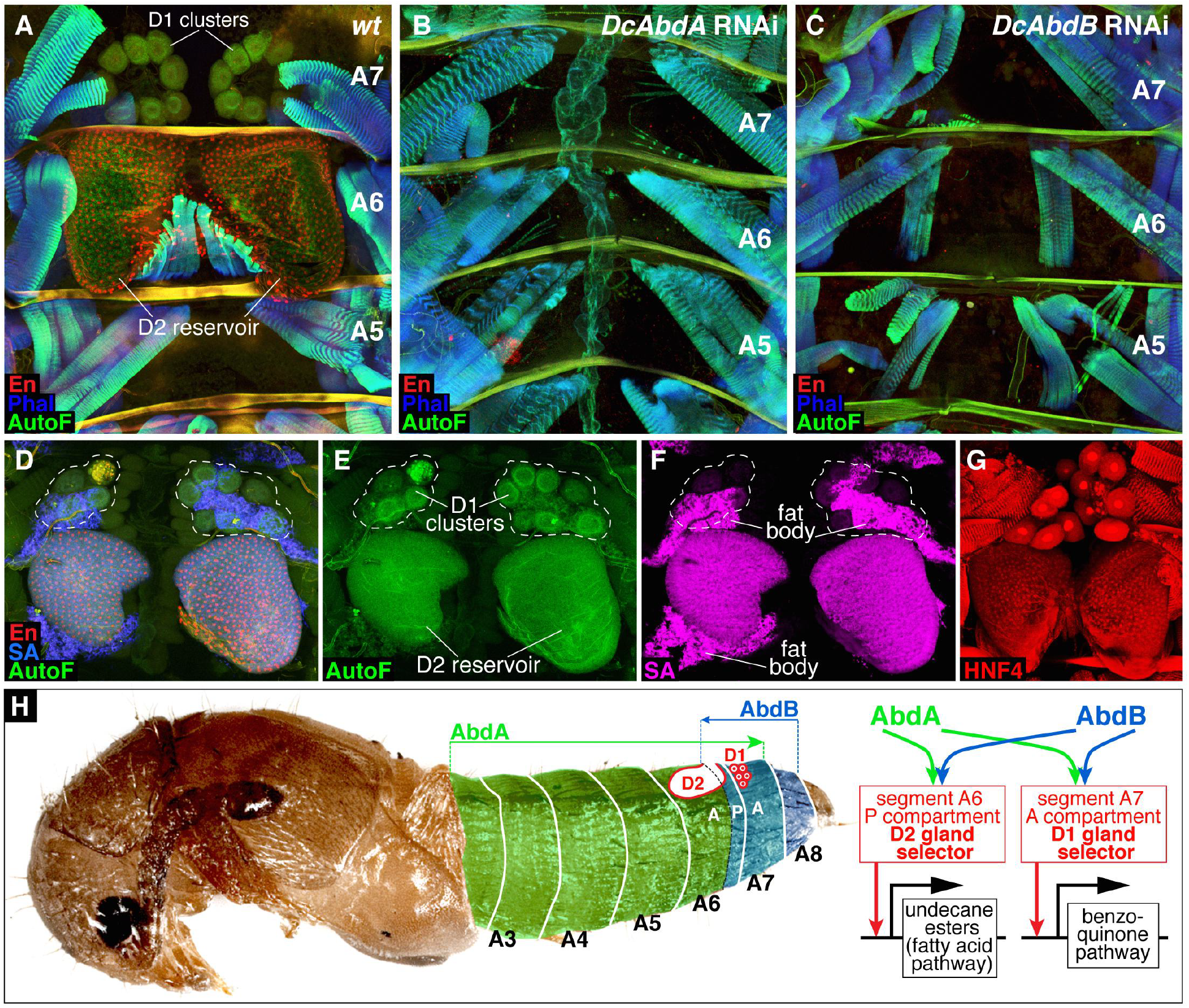
Hox-specification of distinct gland cell types in P and A compartments confers modularity in defensive compound biosynthesis. **A–C**: Confocal images of the dorsal internal adult abdomen of *Dalotia*, labeled for En protein (red), Phalloidin-stained muscle (blue) and autofluorescence (green). **A**: Wild type abdomen. **B**: *DcAbdA* RNAi blocks tergal gland development, with D1 gland units and D2 reservoir missing. **C**: *DcAbdB* RNAi similarly inhibits tergal gland formation. **D–F**: Adult *Dalotia* abdomen labeled for En protein (red), Streptavidin (SA; blue in D, magenta in separated channel in F) and autofluorescence (AutoF; green). The D1 gland unit clusters are indicated with a dashed line in all panels. the D2 cells stain strongly for Streptavidin, but the D1 cells do not (most clear in F). Residual fat body cells attached to the D1 and D2 tissues are also Streptavidin-positive. G: HNF4 stains nuclei of D2 reservoir cells but not D1 gland unit nuclei; note that the red spots in D1 gland units are not nuclei, but non-specific secondary antibody accumulation in the bulbs of D1 gland cells. **H**: Model for *Hox-* logic of tergal gland development and defensive compound biosynthesis. Domains of influence of proteins DcAbdA and DcAbdB are indicated, with a region of overlap that includes the P compartment of segment A6 and A compartment of segment A7. We posit that DcAbdA and DcAbdB act combinatorially to specify different gland cell types in the A6 P compartment and A7 A compartment by acting via distinct intermediate “biosynthetic selector” proteins—transcription factors that govern expression of batteries of different downstream enzymes. In the A6 P compartment DcAbdA+AbdB act via a D2 gland cell selector to specify the D2 reservoir invagination, and expression in D2 cells of enzymes controlling fatty acid metabolism that permit biosynthesis of the alkane solvent and surfactant esters. In the A7 A compartment, DcAbdA+AbdB act via a D1 gland cell selector to specify D1 gland units, which presumably delaminate from the ectoderm, and their enzymatic capacity to synthesize benzoquinones.

### Compartmentalization and modularity of defensive chemical biosynthesis

As well as being anatomically and developmentally distinct, the D1 and D2 gland cell types have been shown to exhibit major ultrastructural differences in the secretory apparatus through which products are transferred into the reservoir (Araujo & Pasteels, 1985). The two cell types thus likely synthesize distinct compounds that contribute to the final, bioactive defensive secretion (Araujo & Pasteels, 1985; Steidle & Dettner, 1993). The n-undecane is an alkane that derives from fatty acids, which are reduced to aldehydes and then decarbonylated (Steidle & Dettner, 1993); similarly, the esters likely derive from fatty acid esterification. We determined whether either of the two gland cell types show evidence of fatty acid metabolism by using fluorescently-tagged Streptavidin, which binds to biotin, a coenzyme in the synthesis of fatty acids from Acetyl-CoA (Fletcher & Myant, 1960; Wakil, Titchener, & Gibson, 1958). Biotin is also involved in the citric acid cycle, but increased cellular labeling of Streptavidin is indicative of an elevated cellular requirement for biotin, and occurs in cell types involved in fatty acid synthesis such as fat body and oenocytes (Burns, Gutzwiller, Tomoyasu, & Gebelein, 2012). We observed a strong and specific upregulation of Streptavidin staining in the D2 reservoir cells, as well as in fat body cells (Fig 5D–F), indicating high levels of biotin. In contrast, Streptavidin failed to detectably bind to the D1 gland cells above background levels (Fig 5D–F). The D2 cells thus appear to be a major site of fatty acid metabolism, consistent with their governing the biosynthesis of the alkane and ester portions of *Dalotia’s* glandular secretion. This hypothesis is consistent with the ideas of Araujo and Pasteels, who argued based on electron microscopic evidence that the D2 cells of the aleocharine *Drusilla canaliculata* control synthesis of the hydrocarbon solvent (Araujo & Pasteels, 1985). To further verify this notion, we labeled the gland for the transcription factor HNF4, which regulates fatty acid metabolism in both the fat body and oenocytes of insects (Burns et al., 2012; Makki, Cinnamon, & Gould, 2014; Palanker, Tennessen, Lam, & Thummel, 2009). Again, HNF4 antibody labeled the nuclei of the D2 cells and fat body, but was absent from the nuclei of D1 cells (Fig 5G, Fig S4D-I).

We propose that the D2 cells synthesize the solvent and esters by expressing enzymes controlling fatty acid metabolism. In contrast, the benzoquinones are presumably synthesized by the D1 glandular units. Structures closely resembling the D1 units produce quinones in other beetle species (Happ, 1968; Li *et al*., 2013). Although the enzymatic pathway(s) remain unidentified, they can utilize tyrosine or polyketide precursors (Pankewitz & Hilker, 2008), which are processed in the gland bulb before being transported along the duct and solubilized in the alkane solvent (Happ, 1968). Araujo and Pasteels (1985) presented support for an analogous scenario in *Drusilla*, and Steidle and Dettner for a variety of other aleocharines (Steidle & Dettner, 1993). We thus think it likely that the purpose of the D1 glandular units is to contribute the benzoquinone fraction of *Dalotia’s* defensive secretion.

## Discussion

Perhaps no other animal group has been so predisposed to evolve complex interspecies relationships as have the aleocharine rove beetles. The dramatic behavioral, chemical and morphological adaptations of many socially parasitic aleocharine species are reflected in associations with ants and termites that rank among the most intimate symbioses known in the Metazoa (Kistner, 1979; 1982; Parker, 2016; Seevers, 1957; 1965). The repeated evolution of symbiosis has its putative basis in preadaptations that the vast majority of free-living species possess, which predispose these beetles to engage in ecological interactions with social insects (Parker, 2016). Chief among these preadaptations are the short elytra that expose the abdomen, and the concomitant presence of a large, targetable defensive tergal gland. By studying *Hox* functions in the free-living aleocharine *Dalotia*, we have uncovered the developmental basis for how this novel preadaptive groundplan evolved within the Coleoptera. These developmental insights provide a foundation for understanding how the anatomy and chemistry of aleocharines have undergone further modification in socially symbiotic species.

### Hox-logic of the preadaptive aleocharine groundplan

We suggest an evolutionary sequence in which Staphylinidae with short elytra and exposed abdomens first evolved from ancestral beetles with long elytra (Fig 6A). This step involved evolutionary changes in the Hox-free ground state circuitry in T2, which reduced the size of the elytron, together with corresponding regulatory changes downstream of Ubx that blocked this size reduction from happening in the T3 hind wings (Fig 6B). Further evolution of the wing folding mechanism had to occur to facilitate packing of the hind wings underneath the small elytra (Saito *et al*., 2014). An exposed, flexible abdomen that is unhindered by overlying elytra consequently arose in Staphylinidae, without sacrificing the beetle’s capacity for flight. This same, basic body plan organization is shared by the majority of the 63,000 species of rove beetle, and was a precondition for the evolution of abdominal chemical defense glands in a variety of subfamilies (Araujo, 1978; Dettner, 1993; Thayer, 2005). In the 16,000 species of Aleocharinae, chemical defensive capacity was accomplished through the evolutionary development of the tergal gland, which arose along the higher aleocharine stem lineage (Fig 6B). This morphological and chemical innovation originated through the synergistic action of the abdominal Hox proteins, AbdA and AbdB, in abutting P and A compartments of segments A6 and A7 where they specify distinct D1 and D2 glandular structures. These glands appear to synthesize distinct classes of compounds, which become bioactive when mixed in the reservoir, leading to total gland functionality (Fig 5H). We note that partitioning biosynthetic processes into different gland cell types, all of which feed into a reservoir, enables aleocharines to create a defensive cocktail from compounds that would, in isolation, be of limited functionality. The alkane and esters are not effective deterrents relative to benzoquines (Steidle & Dettner, 1993), and the benzoquinones would solidify without the alkane solvent. The gland thus displays emergent functionality, whereby the actions of distinct cell types synergize to perform a task not possible by an individual component (Rajapakse & Smale, 2017). This biosynthetic partitioning across cell types is programmed ultimately by AbdA and AbdB acting together but via different outputs in abutting P and A compartments (Fig 5H). By producing distinct gland cell types, a modularity in chemical biosynthesis is achieved that likely facilitates the evolvability of tergal gland secretions, reflected in the diversity of compounds that different species can produce (Stoeffler et al., 2007; 2011; 2013).

**Figure 6.**
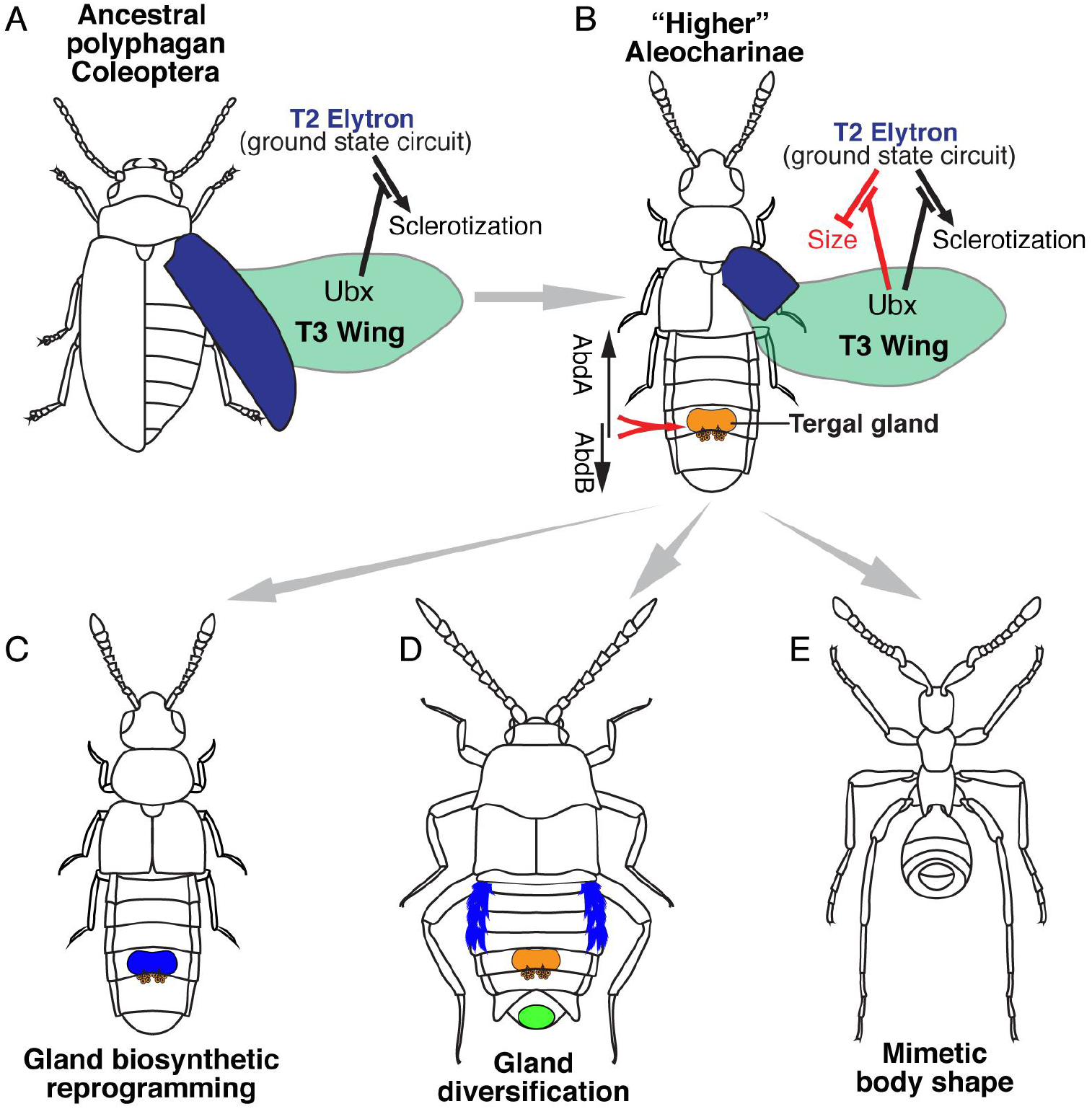
Novel *Hox* functions in the evolution of the aleocharine body plan. **A**: Aleocharine staphylinids evolved within the Coleoptera suborder Polyphaga, from ancestors with long elytra (the scirtid *Cyphon*, a basal polyphagan, is depicted). The beetle elytron (blue) develops in the mesothoracic segment (T2) without *Hox* input, and is heavily sclerotized compared to the membranous forewings of other insects. In T3, *Ubx* overrides both the beetle-specific elytron program, creating membranous wings (Tomoyasu et al., 2005). In staphylinids (B), the elytron circuit has been further modified so the appendage attains only a small size, giving the family’s trademark short elytra; concomitant changes in target genes downstream of *Ubx* in T3 block both the Coleoptera-specific sclerotization program and staphylinid-specific size reduction program, creating enlarged, flight-capable hind wings. Short elytral expose the abdomen, and in higher Aleocharinae, a quinone-based defensive tergal gland (yellow) is specified by the two abdominal Hox proteins, AbdA and AbdB, acting combinatorially in segments A6 and A7. **C–E**: The higher aleocharine body plan, with targetable chemical defense capacity, confers efficient protection from ants and termites, promoting facultative exploitation of colonies. Fully symbiotic species have further modified this preadaptive groundplan by: (C) reprogramming tergal gland biosynthesis to produce compounds for host behavioral manipulation; (D) adding further novel, targetable glands in new abdominal positions for host manipulation; (E) evolving host-mimicking body shapes, in large part by developmental remodeling of the exposed abdomen.

To generate both the D1 and D2 gland types, we propose that AbdA and AbdB were recruited to combinatorially induce expression of secondary transcription factors that execute distinct programs of gland cell specification in A and P compartments (Fig 5H). These putative “biosynthetic selectors” are posited to govern the morphogenesis and differentiation of each of the D1 and D2 tergal gland cell types, as well as these cells’ capacities to synthesize different compounds (Fig 5H). Such a scenario is analogous to neuron type specification, where different “terminal selector” transcription factors have been shown to control neuronal properties by driving expression of batteries of target genes involved in neurotransmitter synthesis, secretion and reception (Hobert, 2011). The identities of the putative biosynthetic selectors are currently unknown, but we note that the invagination of the D2 reservoir is remarkably imaginal disk-like (Fig 4C–E). Appendage patterning genes may thus have been coopted for D2 development, much as they have been for other novel appendage-like structures in other beetle taxa, such as fighting horns (Moczek & Rose, 2009). However, RNAi knockdown of two genes with prominent roles in appendage formation in insects, *Distalless (Dll)* and *Vestigial* (Vg), did not affect tergal gland formation (Fig S6C, D, F) despite inducing the expected appendage phenotypes (Fig S6A, E). Evidently at least these two transcription factors are not gland selector proteins.

We also note that the D2 gland cells share some properties with oenocytes—the abdominal cell type responsible for cuticular hydrocarbon biosynthesis in insects (Gutierrez, Wiggins, Fielding, & Gould, 2006; Makki et al., 2014). Both the D2 cells and oenocytes are ductless gland cells that originate within the P compartment, and their specification depends on AbdA (Brodu, Elstob, & Gould, 2002; 2004). D2 cells and oenocytes are also sites of fatty acid metabolism, with both cell types expressing HNF4 and labeling strongly for Streptavidin (Burns et al., 2012; Palanker et al., 2009). We think it possible that partial recruitment of the oenocyte program, by AbdA and AbdB and putative downstream biosynthetic selectors, may have been involved in D2 cell type evolution. The additional requirement for AbdB in D2 specification could explain why the reservoir only forms in the P compartment of segment A6 where we infer AbdA and AbdB are co-expressed (Fig 4F), whereas oenocytes occur in P compartments of most abdominal segments that lack AbdB expression (Makki et al., 2014). A major enzymatic difference between D2 cells and oenocytes would be that the fatty acid pathway in the D2 cells must lack activity of the elongase necessary to produce a variety of long chain cuticular hydrocarbons (Wicker-Thomas & Chertemps, 2009), leading to production of only short chain alkanes and esters. Further studies of the transcriptomes of the D1 and D2 cells are needed to identify the factors downstream of AbdA and AbdB that govern the differentiation of these gland cell types and their distinct biosynthetic capabilities. We also cannot presently rule out potential microbial involvement in some key steps in the synthesis of tergal gland secretions—for example, the production of the aromatic ring of the benzoquinones is a covalent modification understood in bacteria but not in animals (Pankewitz & Hilker, 2008).

### Modification of the preadaptive groundplan in symbiont lineages

Building on the developmental changes that established the preadaptive groundplan of the higher Aleocharinae, multiple lineages have evolved to specialize on ant and termite colonies, adapting in a finite number of ways (Parker, 2016) (Fig 6C–E). One of these ways is by modifying tergal gland chemistry to produce new compounds that more potently affect host behavior (Fig 6C) (Stoeffler et al., 2007; 2011; 2013). The genetic architecture of the tergal gland hints at how this “chemical reprogramming” could be achieved. Changes in transcriptional targets of a putative biosynthetic selector could lead to recruitment of a new enzyme or enzymatic network into either the D1 or D2 gland cells (or, conversely, loss of expression of one or more biosynthetic enzymes). Across the Aleocharinae, the same basic tergal gland chemistry, comprised of quinones and hydrocarbon solvent, is relatively invariant, suggesting a core biosynthetic apparatus that is conserved in the majority of the subfamily (Steidle & Dettner, 1993). However, the specific benzoquinones and hydrocarbons can differ markedly between species (Steidle & Dettner, 1993) implying species-specific refinement of tergal gland chemistry, presumably via changes in biosynthetic pathways that confer subtle covalent modifications to different compounds within the defensive mixture. The ability of symbiotic species of *Pella* to synthesize the host ant alarm pheromone sulcatone, in addition to undecane and benzoquinones (Stoeffler et al., 2007; 2011), may reflect the co-option of a single terpene pathway into the tergal gland. In contrast, in species of *Zyras*, there has been a wholesale replacement of the typical hydrocarbon-quinone secretion with multiple terpenes that may mimic ant-tended aphids (Stoeffler *et al*., 2013). In this case a more dramatic reprogramming of the underlying biosynthesis must be invoked.

In addition to modifying tergal gland secretions, many symbiotic aleocharine taxa have evolved additional, novel glands besides the tergal gland, typically in other abdominal segments (Fig 6D) (Hölldobler, 1970; Hölldobler, Möglich, & Maschwitz, 1981; Jordan, 1913; Kistner, 1979; 1993; Parker, 2016; Pasteels, 1968; 1969). The compounds produced by any of these glands are unknown, but behavioral observations in a variety of species indicate they function to manipulate or appease host ants and termites. Having already evolved the capacity to develop a complex tergal gland on the abdomen, it is possible that some of these novel glands arose via developmental redeployment of parts of the tergal gland circuitry in new abdominal positions along the anteroposterior axis, presumably under the control of the AbdA and AbdB Hox proteins (i.e., analogous to tergal gland formation, but contingent on a different positional Hox code). Along similar lines, we have previously suggested that the segmentally repeated, paired glands of genera such as *Lomechusa* that promote adoption of the beetle into ant colonies (Hölldobler, 1970)may be modified oenocytes (Fig 1C). Hence, abdominal glands with novel functions in symbiosis could arise through cooption or modification of preexisting glands, as well as via entirely new circuitry. Regardless of how new glandular functions have evolved, however, the fundamental anatomical change that facilitated chemical innovation in aleocharines is the exposed abdomen, which is conducive to gland evolution because it is not masked by elytra, and so glands open directly onto the surface of the beetle. Consequently, the aleocharine abdomen has become an important interface between the beetle and its hosts—an interface shut off to most other Coleoptera which possess long elytra. This same preadaptation of short elytra and abdominal exposure surely underlies the widespread remodeling in symbiont species of abdominal shape, into forms that mimic host ant and termites (Fig 6C) (Kistner, 1979; Maruyama & Parker, 2017; Parker, 2016; Seevers, 1957; 1965). Like the establishment of abdominal glands, evolutionary changes in the sizes and shapes of abdominal segments have presumably been achieved through abdominal Hox-modulation of segment and compartment growth along the anteroposterior axis (Fig 1D, E).

While numerous arthropod lineages have evolved to specialize on social insect colonies as a resource, few have done so as effectively and repeatedly as Aleocharinae. Consequently, the subfamily is a unique system for understanding the evolution of symbiotic interactions between animals. We have invoked a preadaptive anatomical and chemical groundplan of the higher Aleocharinae as having been central to the rampant, convergent evolution of social symbiosis in this clade. We have studied the initial establishment of the groundplan as a way to understand the possible evolutionary starting conditions for symbiosis. Future studies on the molecular and neurobiological modifications to this groundplan seen in symbiotic taxa may provide a framework for comprehending proximate mechanisms governing metazoan symbioses.

## Acknowledgements

We thank Brian Spencer (Applied Bionomics, Canada) for providing *Dalotia*, and Raymond Cloyd (Kansas State University), Graeme Murphy (Ontario Ministry of Food and Agriculture) and Richard Greatrex (Syngenta Bioline, UK) for advice on rearing this species. Yoshinori Tomoyasu provided important feedback on the manuscript. We are grateful to Gary Struhl (Columbia University) in whose laboratory part of this work was carried out.

## Materials and Methods

### *Dalotia* husbandry

*Dalotia* were donated for this study by Applied Bionomics, Canada. Beetles were kept at room temperature in 3- or 5-cup Rubbermaid boxes with a 0.5-1-inch layer of loose coconut fiber substrate (Eco Earth). The substrate was kept slightly damp by occasional misting of the container. Beetles were fed a 1:1 mixture of oatmeal and poultry-rearing pellets powdered in a coffee grinder. Food was sprinkled into the containers and shaken into the substrate every 2–3 days. Adult beetles reach can high densities in containers and display cannibalism, especially of larvae, so were frequently seeded into new containers to avoid population crashes.

### RNA extraction, RNAseq and transcriptome assembly

Two frozen individuals each of the adult, pupal, and larval stages were combined for total RNA extraction with Trizol™ reagent. Library preparation followed the Illumina TruSeq mRNA stranded kit protocol, including chemical shearing to obtain an average final library size of 300 bp. Illumina 100 bp paired-end sequencing was performed on a HiSeq2000 platform in which the *Dalotia* library was multiplexed with eight other samples in one lane. *De novo* transcriptome assembly was performed using Trinity v2.1.1 (Grabherr *et al*., 2011).

### Identification and phylogenetic analysis of *Dalotia Hox* genes

We used tBLASTN to search the *Dalotia* transcriptome for *Scr, Antp, Ubx, AbdA* and *AbdB* using protein sequences of their *Tribolium* homologs. We recovered clear single orthologs of each protein, and aligned these with sequences from the beetles *Dendroctonus* and *Nicrophorus* and *Tribolium*, and added further ortholog sequences from *Drosophila* and *Apis*. The prototypical Hox protein from *Trichoplax adherans*, Trox-2, (Jakob *et al*., 2004) was included as an outgroup. Bayesian analysis was performed in MrBayes 3.2 (Ronquist *et al*., 2012)

### DsRNA preparation and RNAi knockdown in *Dalotia*

DsRNA was prepared from mixed cDNA from pooled larval and adult animals, and regions of 300-600 bp were amplified using primers with T7 linkers. Fragments were cloned into pCR4-TOPO (Thermofisher). The following primers were used:

DcWhite-RNAiF1 TAA TAC GAC TCA CTA TAG GGC GTC CGG GTG AAT TGT TAG C

DcWhite-RNAiR1 TAA TAC GAC TCA CTA TAG GGT CAG CAG GGT TGT AAT TAT GCG

DcVer-RNAiF1 TAA TAC GAC TCA CTA TAG GGG AGA ATG CTC AGT GGC AAC G

DcVer-RNAiR1 TAA TAC GAC TCA CTA TAG GGC ACC GCG TTT AAC CAA TGC C

DcScR-RNAiF1 TAA TAC GAC TCA CTA TAG GGC AGC ATC CCT ATG CAA CAC C

DcScR-RNAiR1 TAA TAC GAC TCA CTA TAG GGA CCA TTC GCA TTC ACT GTA CG

DcAntP-RNAiF2 TAA TAC GAC TCA CTA TAG GGG ATC CGA ATT CCT GTG GTG G

DcAntP-RNAiR2 TAA TAC GAC TCA CTA TAG GGG GAA CTC CTT CTC TAG CTC C

DcUbxRNAiF1 TAA TAC GAC TCA CTA TAG GGC CTT CTT CAG CTT CAT GCG G

DcUbxRNAiR1 TAA TAC GAC TCA CTA TAG GGC GGA GTA AAA GAC GTG TGG C

DcAbdA-RNAiF1 TAA TAC GAC TCA CTA TAG GGC CAA ATT CAT CAT CGA TAG CAT G

DcAbdA-RNAiR1 TAA TAC GAC TCA CTA TAG GGC TGG GAG CAG GAA TTC AAC G

DcAbdB-RNAiF2 TAA TAC GAC TCA CTA TAG GGT CTC ATC GTG CCA TCA GAG C

DcAbdB-RNAiR2 TAA TAC GAC TCA CTA TAG GGG TGA TGA TTA ACA ACG TGG TGC

DcApA-RNAiF1 TAA TAC GAC TCA CTA TAG GGC GTC TAA CAA CAA TCA CTT GGC

DcApA-RNAiR1 TAA TAC GAC TCA CTA TAG GGC TAC CGG TGG TAT AGA TGA CG

DcApB-RNAiF1 TAA TAC GAC TCA CTA TAG GGC GAT CGC TAC TAT CTG CTT GC

DcApB-RNAiR1 TAA TAC GAC TCA CTA TAG GGG CAT CTG GAT TCT GAT TGA TAG C

DcDll-RNAiF1 TAA TAC GAC TCA CTA TAG GGC TCA AAG TCG GCG TTC ATC G

DcDll-RNAiR1 TAA TAC GAC TCA CTA TAG GGC CTC CTT GCA TCA TAT TCT GG

DcVg-RNAiF1 TAA TAC GAC TCA CTA TAG GGG TGC TGC ACA TCA ATA TGA TAG G

DcVg-RNAiR1 TAA TAC GAC TCA CTA TAG GGG GTG ATG GTA GTC GTG AAC G

The same primers were subsequently used to make template DNA strands for DsRNA synthesis, by amplifying the fragments that were inserted into the TOPO vector and using the Megascript T7 high yield transcription kit (Thermofisher). DsRNA was typically diluted to a 2× concentration of 4 mg/ml, and then diluted 1:1 in PBS and green food dye following a previously published protocol (Philip & Tomoyasu, 2011). For microinjections, injection needles were made from capillary tubes using a micropipette puller. Each needle was back-loaded with 2-3 μL of dsRNA solution and then attached to a syringe. *Dalotia* specimens were collected and injected at the late larval stage. Late third instar larvae were collected from populations using an aspirator, and 5-7 animals were placed on a CO_2_ gas pad and mounted using a paintbrush onto a strip of double-sided tape on a microscope slide. The slide consisted of two microscope slides attached with double-sided tape, misaligned to create a 1 cm edge. During the injection process, CO2 was blown over the slide to keep the larvae anaesthetized. Each larva was injected with ~0.5 μL dsRNA solution, until the larval body appeared slightly swollen and green in color due to the food dye. After injection, the larvae were allowed to awaken and walk off the tape. The larvae were then placed into moistened scintillation tubes containing plaster of Paris, in which most animals pupated within 24–48 hours. After ten days, adults or uneclosed pupae were inspected for the mutant phenotype. Larvae that died before pupation or did not pupate by the end of ten days were disregarded.

### Immunohistochemistry

Adult beetles were immersed in PBS and abdomens were removed with forceps. The ventral segments were removed by cutting around the abdominal margin with dissection scissors. The dorsal abdomen was fixed in 4% paraformaldehyde (25 minutes, room temperature), washed in PBS+0.02% Triton X100, and stained with the following antibodies: anti-Engrailed 4D9 (1:5; DSHB), anti-*Tribolium* HNF4 (1:500; B. Gebelein), Anti-AbdA/Ubx FP6.87 (1:10; DSHB). Alexa-Fluor secondary antibodies (Thermofisher) were used, along with Alexa-647-Phalloidin (Thermofisher) to label muscles, Alexa-647-Streptavidin (Thermofisher) to reveal biotin, and Hoechst to mark nuclei. Tergal glands were imaged in whole mount dorsal abdomens in Vectashield (Vectorlabs), using a Leica SP5 microscope.

### Gland Histology

Pupae were staged to 24-hour intervals, freshly killed and fixed in 24% glutaraldehyde for 1 day. Specimens were then dehydrated through a dilution series to 100% EtOH. Infiltration into LR White epoxy proceeded through a dilution series of 100% EtOH:LR White mixture, allowing tissues to incubate in pure LR White for ~12 hours. Embedding proceeded by placing specimens in gelatin capsules filled to the top with LR White and then in an oven for 24 hours at 60 °C (thermal curing). Embedded specimens were sectioned using a Leica EM UC6 ultramicrotome and diamond knife, producing sections 5-6 μm thick. Sections were stained with toluidine blue. Specimens embedded in Spurr’s Resin were dehydrated in an Acetone series prior to infiltration and embedding. Sections were imaged on a Zeiss Axioplan 2 microscope.

### Construction of *UAS-DcUbx* and *UAS-DmUbx*

A full length *DcUbx* coding sequence was amplified from *Dalotia* larval and adult mixed cDNA using primers designed using the *DcUbx* sequence recovered from the transcriptome. A full length *DmUbx* coding sequence was also obtained (gift from R. Mann). A 3xHA tag was placed at the N-terminus of both coding sequences, and the constructs were ligated into pUAS-ATTB (Bischof, Maeda, Hediger, Karch, & Basler, 2007) using KpnI and XbaI linkers. *UAS-3xHA-DcUbx* and *UAS-3xHA-DmUbx* were then inserted into the same chromosomal location in *Drosophila*, at 27C2 using phiC31 integrase-mediated recombination (Bischof *et al*., 2007).

### *Drosophila* genetics

A stock containing *Nubbin-GAL4* and *GAL80^ts^* transgenes was crossed to *UAS-DcUbx* or *UAS-DmUbx* flies. After a 24 hour collection, larvae were transferred to 18°C and maintained at low densities to prevent crowding affecting wing size. At the start of the third instar, larvae were transferred to 29°C to permit GAL4-driven *Ubx* expression in the wing blade. Adult wings were dissected and mounted in Hoyer’s medium, and imaged with a Zeiss compound microscope. To assess the affects of to *UAS-DcUbx* or *UAS-DmUbx* on Dpp production, *dpp-GAL4* was used to drive *UAS-Ubx* transgenes in the Dpp expression domain of the wing imaginal disc. Late third instar discs were dissected and fixed in 4% paraformaldehyde, and guinea pig phospho-MAD antibody (1:1000; E. Laufer) was used to assess the range of Dpp transduction.

### Gas chromatography/Mass spectroscopy of tergal gland contents

Three methods were used to investigate the composition of the tergal gland secretion:

i. Submersion of the beetles in hexane for 1 minute: gland exudate extraction was accomplished by submersing four living individuals in 50 μl hexane; after 1 minutes the solvent was separated from the beetles.
ii. Pressing filter paper against the gland opening and subsequent extraction of the filter paper in 50 μl hexane. Additionally, filter paper without gland exudates was extracted as a control.
iii. Dynamic headspace sampling of the beetle gland secretions. Groups of four beetles were placed in a glass vial and sealed with a lid. The lid was pierced with a glass tube (13mm; ID 5mm) filled with a mixture (1:1) of 3 mg Tenax-TA (mesh 60–80; Supelco, Bellefonte, Pennsylvania, USA) and 3 mg Carbotrap B (mesh 20–40, Supelco, Bellefonte, Pennsylvania, USA), and used as adsorbent tubes. The adsorbent was fixed in the tubes with glass wool. To stimulate the release of gland secretion, beetles were gently stirred with a small magnetic stick on a magnetic stirrer for 20 minutes and the headspace was continuously sampled using a Tuff™ pump system (Casella, Bedford, UK) with a flow rate of 80 ml/min. Simultaneous collection of the surrounding air was performed to distinguish ambient contaminants. Afterwards, adsorbent tubes were extracted with 50μl hexane.

A GCMS-QP2010 Ultra gas-chromatography mass-spectrometry system (Shimadzu, Duisburg, Germany) equipped with a ZB-5MS fused silica capillary column (30 m × 0.25 mm ID, df= 0.25 μm) from Phenomenex (Aschaffenburg, Germany). Crude hexane sample aliquots (2 to 5 μl) were injected by using an AOC-20í autosampler system from Shimadzu, Duisburg, Germany into a PTV-split/splitless-injector (optic 4, ATAS GL, Eindhoven, Netherlands), which operated in splitless-mode. Injection-temperature was programmed from an initial 40°C up to 230°C and then an isothermal hold for 13 minutes. Hydrogen was used as the carrier-gas with a constant flow rate of 3.05 ml/min. The chromatographic conditions were as follows: The initial column temperature was 50°C with a 1-minute hold after which the temperature was increased 25°C/min to a final temperature of 300°C and held for 2 minutes. Electron impact ionization spectra were recorded at 70 eV ion source voltage, with a scan rate of 0.3 scans/sec from m/z 40 to 400. The ion source of the mass spectrometer and the transfer line were kept at 250°C. Compounds were identified based on their *m/z* fragmentation patters. Additionally, the identity was confirmed by comparison of retention indices and MS data with published literature (Dettner 1984, Dettner et al. 1985, Steidle and Dettner 1993), additionally library comparisons with Wiley Registry of Mass Spectral Data 2009 and NIST 2011 were performed.

## Supplemental Figures

**Figure S1.**
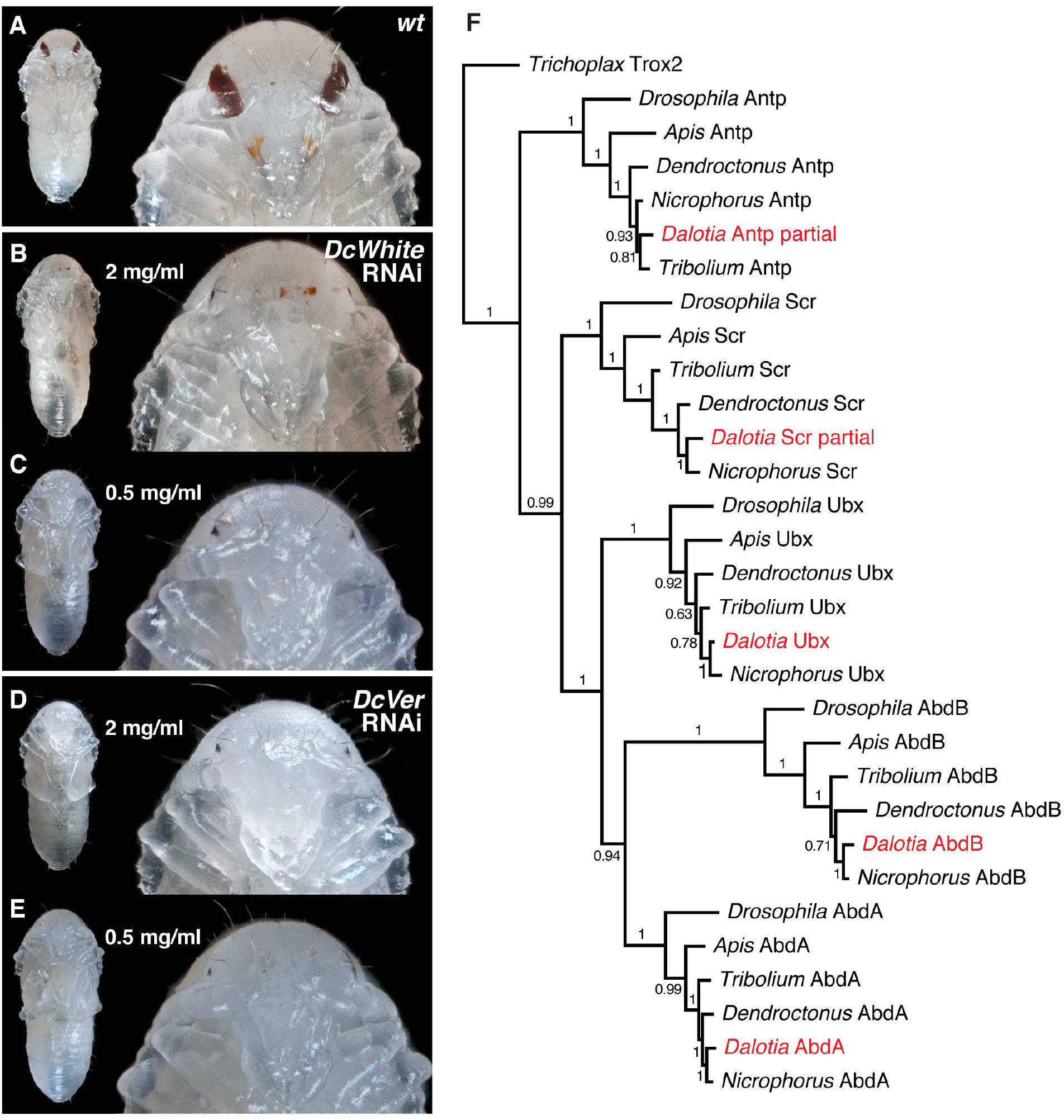
Systemic larval RNAi in *Dalotia*, and the rove beetle trunk *Hox* gene complement. **A**: Wild type pupa showing dark eye pigmentation. **B, C**: DsRNA knockdown of the *Dalotia* white ortholog at 2 mg/ml and (B) and 0.5 mg/ml (C) removes eye pigmentation. **D, E**: DsRNA knockdown of the *Dalotia* vermillion ortholog at 2 mg/ml and (D) and 0.5 mg/ml (E) removes eye pigmentation. **F**: Bayesian phylogenetic tree of Hox protein sequences from *Dalotia* and orthologs from other insect species. The tree is rooted with *Trichoplax* Trox2, a putative sister to the paralogous Hox proteins of eumetazoans.

**Figure S2.**
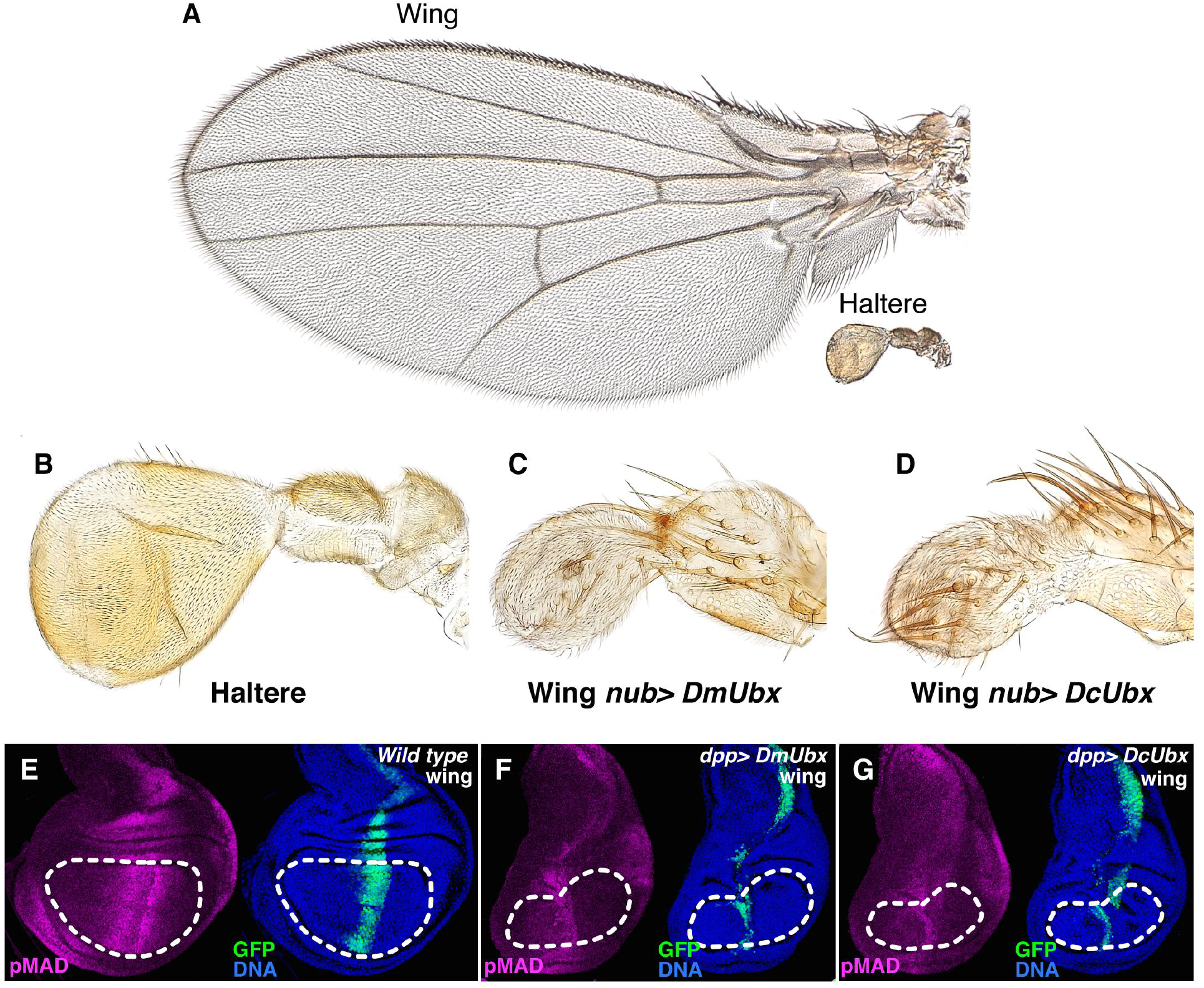
Misexpression of *Dalotia Ubx* in *Drosophila*. **A**: *Drosophila* wing and haltere, to scale, to demonstrate relative sizes. **B**: *Drosophila* haltere, enlarged. **C**: Expression in the wing of either *Drosophila Ubx* (C) or *Dalotia Ubx* (D) reduces wing growth and transforms the organ into a haltere-like structure (B–D are to scale). **E–G**: *Drosophila* wing imaginal discs expressing UAS-GFP (green) with *Dpp-GAL4* and stained with phospho-MAD (pMAD) antibody (magenta) and Hoeschst-labelled nuclei (blue). In otherwise wild type discs (E), a pMAD gradient extends along the anteroposterior axis from GFP-labelled Dpp expression domain. Expression of either *Drosophila Ubx* (F) or *Dalotia Ubx* (G) strongly impedes the range of the pMAD gradient, indicating reduced Dpp spreading. The discs in F and G show a corresponding growth defect.

**Figure S3.**
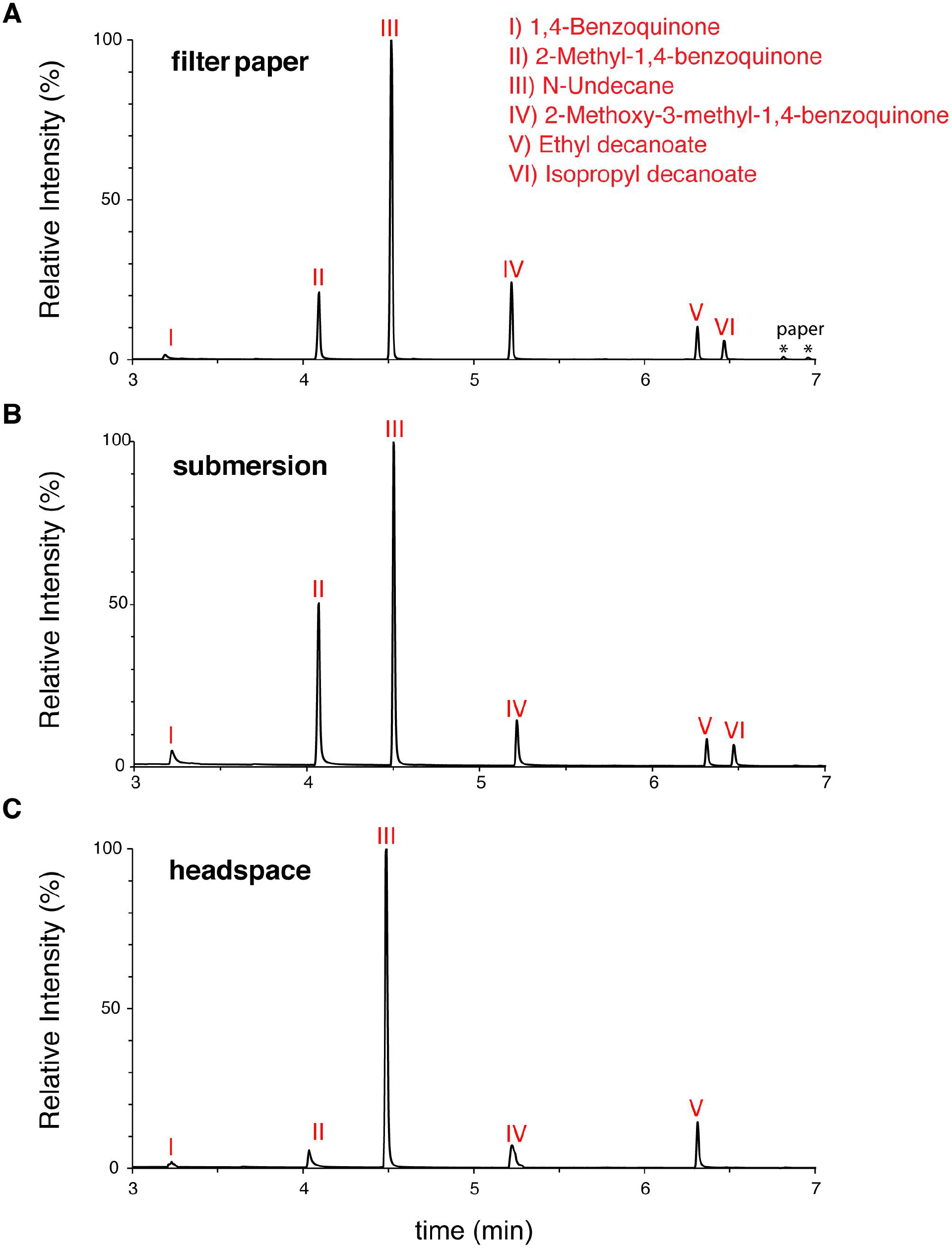
GC/MS of *Dalotia* tergal gland contents. **A–C**: Traces of gland contents assayed by dabbing filter paper the gland opening (A; asterisks are two peaks from the paper), submerging whole beetles briefly in hexane to eject gland contents (B) and sampling headspace volatiles (C).

**Figure S4.**
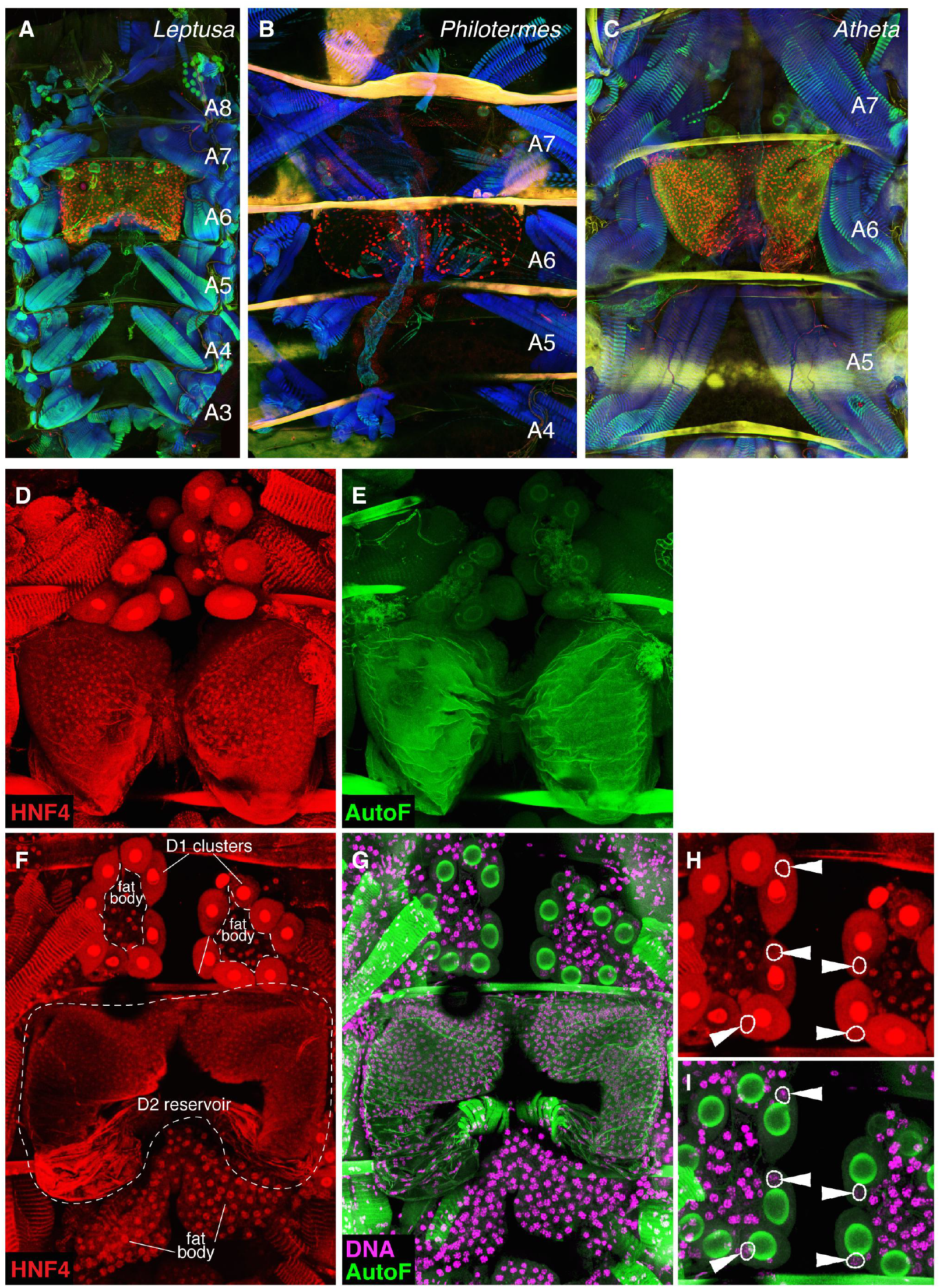
Tergal glands in higher Aleocharinae genera and HNF4 expression in D2 gland cells. **A–C**: Tergal glands in three aleocharines labelled for En protein (red), phalloidin-stained muscle (blue) and autofluorescence (green): *Leptusa* (A), *Philotermes* (B) and *Atheta* (C). **D–I**: HNF4 staining of the *Dalotia* tergal gland. D and E show HNF4 protein (red) and cuticular autofluorescence (green). The D2 gland cell nuclei stain positively for HN4, while the D1 cell nuclei do not. The red spots in D1 glands in panel D are an artifact of antibody accumulation in the bulbs of the D1 cells. F-I show a further example of HNF4-positive D2 cell nuclei and lack of HNF4 staining in nuclei of D1 gland cells, in this case nuclei are counterstained with Hoechst (magenta). A close up of a portion of the gland is shown in panels H and I, with nuclei of D2 gland cells circled

**Figure S5.**
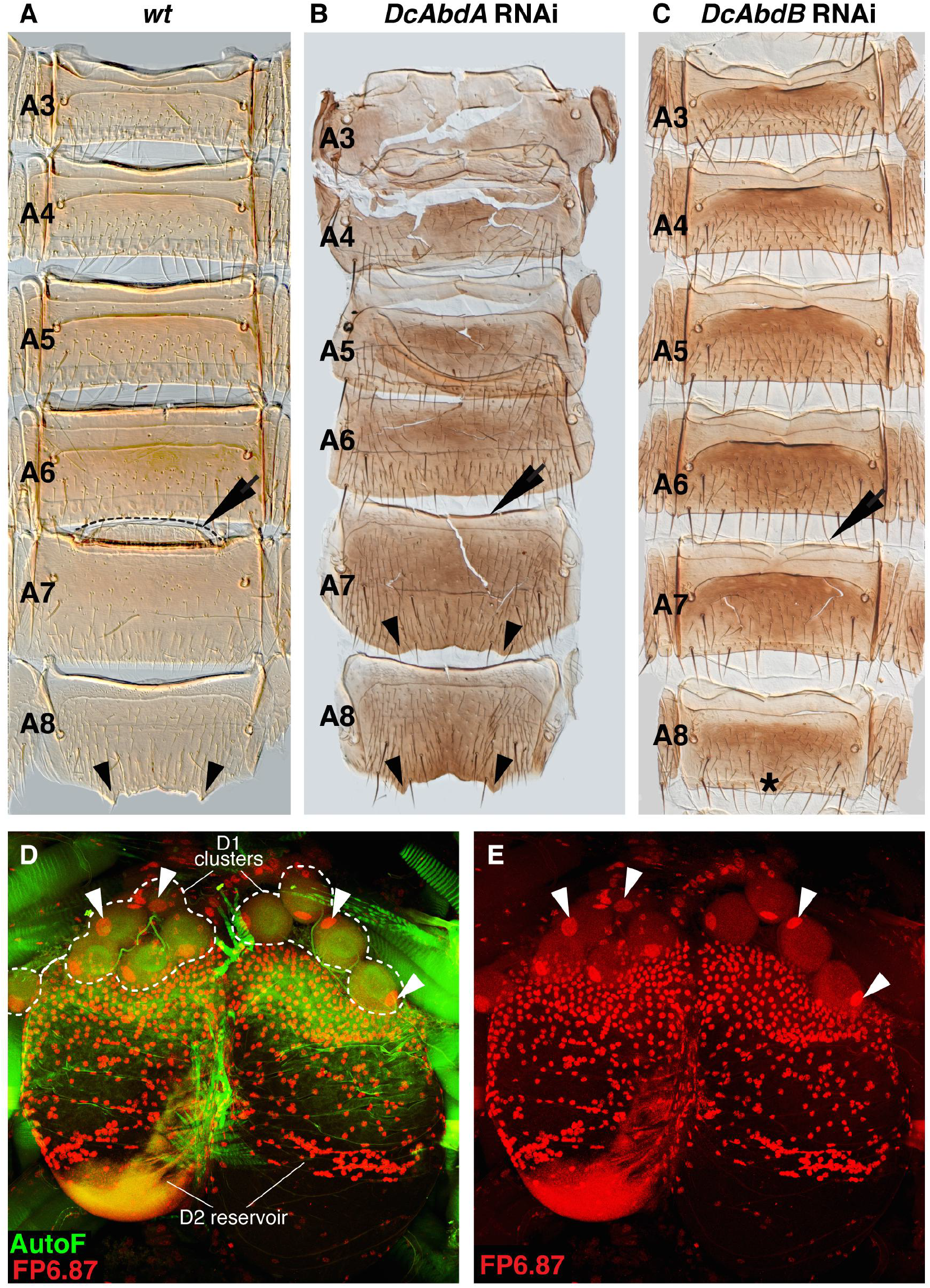
Effects of AbdA and AbdB knockdown on dorsal abdomen cuticle morphology in *Dalotia*. **A**: Male wild type abdomen with shelf of gland opening at the anterior margin of A7 indicated (arrow), and jagged posterior margin of A8 (arrowheads). **B**: AbdA RNAi leads to segmental transformations that include loss of the A7 tergal gland shelf (arrow), and appearance of A8-like morphology in A7. **C**: AbdB RNAi leads to segmental transformations that include loss of the A7 tergal gland shelf (arrow), and loss of the jagged posterior margin of A8 (asterisk). **D, E**: FP6.87 staining (red) labels AbdA protein in nuclei of both D2 and D1 cells (arrowheads point to D1 cell nuclei). Green: autofluorescence. Note that FP6.87 antibody also recognizes Ubx protein.

**Figure S6.**
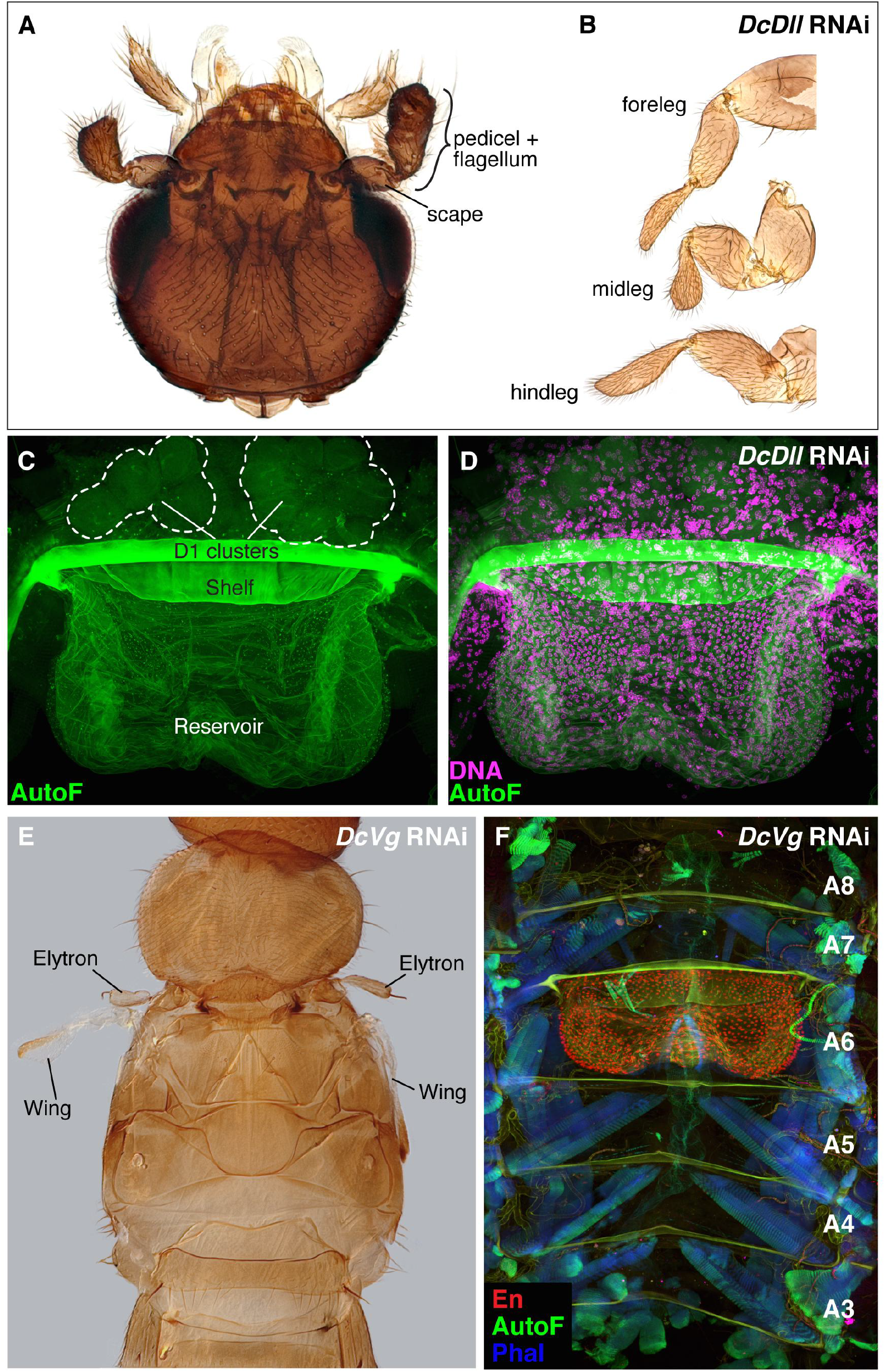
Effects of Dll and Vg knockdown in *Dalotia*. **A–D**: Knockdown of Dll causes severe truncation of the antennae (A) and legs (B), which lose their tarsi. However, the tergal gland is unaffected in these animals (C, D; green: autofluorescence; magenta: nuclei). **E, F**: Knockdown of Vg causes substantial loss of elytron and wing blade structures aside from putative proximal hinge tissue (E), but tergal gland anatomy is unaffected (F; red: En protein, blue: phalloidin-stained muscle, green: autofluorescence).

